# Advancing the Conceptualization and Measurement of Mentorship: The Mentoring Experiences in Research & Graduate Education (MERGE) Scale

**DOI:** 10.64898/2026.09.19.752913

**Authors:** Trevor T. Tuma, Melissa E. Robertson, Erin L. Dolan

## Abstract

Interest in mentoring remains widespread within the scientific community, as mentorship is considered crucial to developing the scientific workforce. A longstanding focus on the positive aspects of mentorship obscures the reality that some mentoring relationships may be ineffective, counterproductive, or harmful. Progress has been limited by the absence of a scale capable of measuring both the positive and negative mentoring experiences of doctoral students. We addressed this need by developing a new scale: the Mentoring Experiences in Research and Graduate Education (MERGE). Results of three sequential multi-method studies (total *N* = 1,308) provide evidence for: (a) the content and response process validity of the MERGE; (b) a six-factor structure capturing the range of mentoring experiences; (c) negative mentoring experiences as empirically distinct from the absence of positive mentoring; (d) convergent, discriminant and criterion-related validity of the MERGE, with each factor demonstrating unique patterns of association with outcomes; and (e) the incremental validity of the MERGE in predicting outcomes beyond an existing measure of mentoring competence. We conclude by discussing implications for the use of the scale in doctoral education, including both theory advancement and applied practice.

## INTRODUCTION

Mentorship is a developmental relationship in which a more experienced individual (the mentor) and a less experienced individual (the mentee) work together to support the personal and professional growth, development, and success of both relational partners (NASEM, 2019). High-quality mentoring is associated with numerous behavioral, career, and health-related benefits for both mentees and mentors (Allen et al., 2004; Eby et al., 2008; Eby et al., 2013; Tenenbaum et al., 2001). In STEM doctoral education, mentorship occurs through relationships between doctoral students and their faculty research advisors, where the support that faculty provide can shape students’ educational experiences and career outcomes. The quality of these relationships is widely regarded as one of the most important influences on student success (Sverdlik et al., 2018; Zhao et al., 2007), and effective mentorship is thought to be critical for training the next generation of PhD-level STEM professionals.

Research in STEM education has primarily focused on supportive mentoring functions, particularly career and psychosocial support (Kram, 1988; Noe, 1988). This emphasis provides only a partial understanding of mentoring by overlooking the possibility that these relationships may be dysfunctional or even harmful (Kram, 1985; Scandura, 1998). Like all close relationships, mentoring relationships vary in their quality and effectiveness (Eby et al., 2000; Ragins et al., 2000), and may include elements that are ineffective, dysfunctional, or actively harmful (Eby et al., 2000, 2004; Limeri et al., 2019; Tuma et al., 2021, 2025). Negative mentoring experiences have been reported by both mentees (Eby et al., 2000, 2004; 2010; Scandura, 1998) and mentors (Eby et al., 2008; Eby & McManus, 2004), and can vary in severity and type (Eby et al., 2000; Eby & Allen, 2002). Such experiences are not uncommon, with 25-50% of mentees experiencing some form of relational dysfunction with their mentor (Clark et al., 2000; Eby et al., 2000).

Although research on negative mentoring experiences in student-faculty relationships is underdeveloped (Allen & Eby, 2007; NASEM 2019), evidence suggests that doctoral students’ mentorship experiences vary even within the same institution, graduate program, and lab (Griffin et al., 2022; Maher et al., 2017; Ruud et al., 2018; Tuma et al., 2021). Scholars have proposed that negative experiences may take multiple forms, ranging from mismatches in communication style, personality, or career interests to problematic mentor behaviors such as neglect, incompetence, interpersonal conflict, and boundary violations (Johnson & Huwe, 2002). Graduate students have likewise reported experiencing sabotage, emotional abuse, harassment, manipulation, credit taking, and neglect from faculty mentors (Castelló et al., 2017; Clark et al., 2000; Johnson, 2003; Kalbfleisch, 1997; Tuma et al., 2021). Given that negative mentoring experiences have been associated with a range of adverse outcomes (Eby et al., 2004, 2010), and destructive supervisory relationships impair employee performance, creativity, organizational commitment, and retention (Harris et al., 2007; Porath & Erez, 2009; Porath & Pearson, 2010; Zellars et al., 2002), there is reason to believe that similar detrimental effects may occur among doctoral students.

Measures developed for workplace and undergraduate research mentoring capture some forms of both positive and negative mentoring but were developed in contexts that differ from doctoral education (Eby et al., 2004; Limeri et al., 2024). In contrast, measures used in doctoral education tend to exclusively assess positive aspects of mentoring or narrow aspects of negative experiences, providing incomplete coverage of the range of experiences doctoral students may encounter (Golde & Dore, 2001; Noy & Ray, 2012; Zhao et al., 2007). Moreover, some measures conceptualize negative mentoring primarily as low levels of positive mentoring (Schlosser & Gelso, 2001). However, a mentoring relationship can provide limited career support without involving actively harmful behavior. Thus, treating positive and negative mentoring as opposite ends of a single continuum may obscure meaningful differences between distinct mentoring experiences. Efforts to advance research on mentoring in doctoral education have been constrained by the lack of measurement tools capable of generating valid and reliable scores to understand the full range of mentoring that doctoral students may experience (NASEM, 2019).

To address this need, we conducted a three-phase study to develop a new measure of doctoral students’ mentoring experiences, which we term the <u>M</u>entoring <u>E</u>xperiences in <u>R</u>esearch and <u>G</u>raduate <u>E</u>ducation (MERGE). We present the development of the MERGE along with evidence of its validity as a measure of the mentoring experienced by doctoral students. The development and validation of the MERGE will advance research, practice, and theory. Specifically, it will (a) enable investigation of the antecedents, correlates, and consequences of the range of doctoral students’ mentoring experiences and the evaluation of various mentoring interventions, (b) equip graduate programs and university leaders with a tool to identify, monitor, and improve the quality of mentorship experienced by doctoral students, and (c) advance theoretical understanding of doctoral students’ mentoring experiences by clarifying the dimensionality of mentoring experiences.

## METHODS AND RESULTS

We conducted three sequential, multi-method studies to provide evidence supporting the construct validity of the MERGE as a measure of doctoral students’ mentoring experiences. We provide an overview of the phases, methods, and samples in Figure 1. The research was determined to be exempt by the University of Georgia Institutional Review Board (PROJECT00003604). Participant information for all three phases is reported in Table 1. Students received a gift card for their participation. Data were analyzed using R (R Core Team, 2021) with functions from the readability (Rinker, 2015), EGAnet (Golino & Christensen, 2024), psych (Revelle, 2011), lavaan (Rosseel, 2012), and yhat (Nimon et al., 2025) packages. This study was not preregistered.

**Figure 1.**
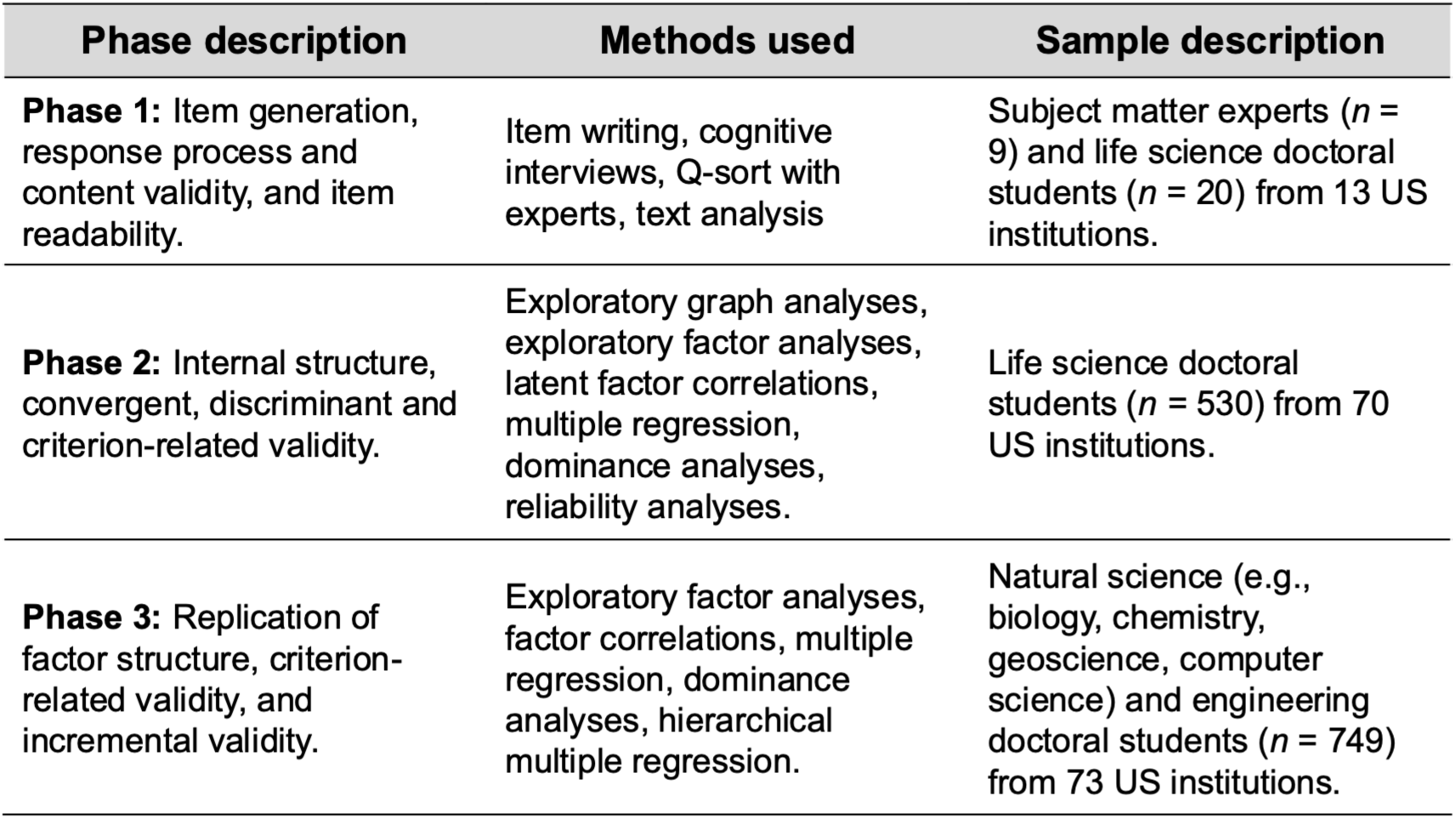
Overview of Study Phases, Methods, and Samples

**Table 1.** Characteristics of Study Participants.

| Description | Phase 1<br>(*n* = 20) | Phase 2<br>(*n* = 530) | Phase 3<br>(*n* = 749) |
| --- | --- | --- | --- |
| **Gender** |  |  |  |
| Woman | 14 (70%) | 306 (58%) | 399 (53%) |
| Man | 6 (30%) | 211 (40%) | 277 (37%) |
| Non-binary | 0 (0%) | 11 (2%) | 21 (3%) |
| Prefer not to respond | 0 (0%) | 2 (<1%) | 49 (7%) |
| **Race\*** |  |  |  |
| American Indian or Alaskan Native | 0 (0%) | 10 (2%) | 4 (1%) |
| Asian | 6 (30%) | 121 (23%) | 198 (26%) |
| Black or African American | 3 (15%) | 49 (9%) | 47 (6%) |
| Hawaiian or Native Pacific Islander | 0 (0%) | 3 (<1%) | 1 (<1%) |
| North African & Middle Eastern | 0 (0%) | 14 (3%) | 24 (3%) |
| White | 10 (50%) | 317 (60%) | 427 (57%) |
| Prefer to self-describe | 0 (0%) | 21 (4%) | 21 (3%) |
| Prefer not to respond | 1 (5%) | 20 (4%) | 47 (6%) |
| **Ethnicity** |  |  |  |
| Hispanic or Latina/Latino | 5 (25%) | 83 (16%) | 78 (10%) |
| Not Hispanic | 15 (75%) | 433 (81%) | 608 (81%) |
| Prefer not to respond | 0 (0%) | 14 (3%) | 63 (8%) |
| **Degree Status** |  |  |  |
| Pre-Candidacy | 7 (35%) | 193 (36%) | 225 (30%) |
| Post-Candidacy | 13 (65%) | 335 (63%) | 477 (64%) |
| Prefer not to respond | 0 (0%) | 2 (<1%) | 47 (6%) |
| **Years Completed** |  |  |  |
| 1 | 2 (10%) | 91 (17%) | 109 (15%) |
| 2 | 4 (20%) | 103 (19%) | 166 (22%) |
| 3 | 4 (20%) | 116 (22%) | 130 (17%) |
| 4 | 4 (20%) | 82 (16%) | 159 (21%) |
| 5 | 2 (10%) | 80 (15%) | 87 (12%) |
| 6 | 4 (20%) | 41 (8%) | 41 (5%) |
| 7+ | 0 (0%) | 13 (2%) | 7 (1%) |
| Prefer not to respond | 0 (0%) | 4 (<1%) | 50 (7%) |
| **Research Context\*** |  |  |  |
| Bench | 13 (65%) | 393 (74%) | 513 (68%) |
| Computational | 10 (50%) | 257 (48%) | 301 (40%) |
| Fieldwork | 5 (25%) | 152 (29%) | 151 (20%) |
| Theoretical | 6 (30%) | 49 (9%) | 72 (10%) |
| Prefer not to respond | 0 (0%) | 1 (<1%) | 48 (6%) |
| **International Student Status** |  |  |  |
| Yes | 6 (30%) | 111 (21%) | 193 (26%) |
| No | 14 (70%) | 418 (79%) | 506 (68%) |
| Prefer not to respond | 0 (0%) | 1 (<1%) | 50 (7%) |

\* = Percentages do not sum to 100% because participants could select multiple response options.

### Phase 1: Item Generation, Response Process, and Content Validity

In Phase 1, we generated items for a scale to measure the range of doctoral students’ mentoring experiences. We then assessed the response process validity and content validity of the item set.

#### Item Generation

To ensure comprehensive coverage of doctoral students’ mentoring experiences, we generated items based on the qualitative themes identified by Tuma and colleagues (2021). The goal of the measure was to capture doctoral students’ perceptions of mentor behaviors and interactions within the mentoring relationship and excluded broader macrosystem (i.e., practices, values, and norms of academic STEM research) and microsystem (i.e., lab, departmental, or institutional factors) factors (Tuma et al., 2021). We began by reviewing the original transcripts, conceptual definitions, and operationalizations derived from doctoral students’ narrative accounts (Tuma et al., 2021). Two of the authors independently drafted items, using students’ own words and aiming to keep statements simple, concise, and free of culture-specific phrases. We then collaboratively reviewed the item pool, removed redundant or overlapping items, and selected items that best represented these experiences while maintaining coverage of the construct. This process resulted in a pool of over 210 items.

#### Response Process Validity

We next examined whether items were interpreted as intended by the target population. Specifically, we evaluated whether doctoral students’ understanding of each item aligned with its intended meaning and whether items elicited their mentoring experiences sufficiently. We conducted semi-structured cognitive interviews using a concurrent think-aloud approach (Willis, 1999), in which participants verbalized their thought processes while responding to items. Participants read and responded to each item, explained their interpretation of the item, and described how they arrived at their response. Given the large item pool and the need to minimize fatigue, each participant reviewed a subset of the items (40 items each); each item was reviewed by at least four doctoral students. We interviewed 20 life science doctoral students representing 13 institutions across the US, with a range of backgrounds and mentorship quality. Participants represented 13 institutions across the US and varied in their English-language status (14 native English speakers, 6 English language learners). Semi-structured interviews were conducted via Zoom and lasted approximately 50 minutes.

Based on interview responses, we revised or removed items that participants identified as ambiguous or unclear. For example, we removed colloquial phrases (e.g., *“passive-aggressive,” “keeps me out of the loop,” “stretch the truth”*) and items that participants deemed too vague or general to gauge (e.g., *“My mentor provides useful feedback,”* “*My mentor lacks the necessary expertise,”* “*My mentor pretends to help me more than they actually do.”*) These revisions resulted in a pool of 103 items.

#### Content Validity

We next examined whether the revised item pool adequately represented the range of mentoring experiences. Nine experts in mentoring, workforce development, industrial-organizational psychology, and survey questionnaire development were recruited to participate in a Q-sort content validation activity to assess the relevance, representativeness, and clarity of the items (Nahm et al., 2002). The experts were provided with a randomized list of items and were asked to evaluate each item’s relevance and appropriateness for measuring the range of students’ mentoring experiences. Experts were also asked to provide qualitative feedback regarding construct coverage (i.e., potential construct underrepresentation or overrepresentation) and identify ambiguous or problematic items. We considered expert feedback about construct coverage, ambiguity, redundancy, and wording to further revise, retain, or remove items. We focused revisions on items with low agreement, as well as those flagged by experts as unclear or misaligned with construct definitions.

We assessed item readability using standard text analysis methods. We examined two well-known and widely used readability indices: the Flesch reading ease score (Flesch, 1948), which reflects how easy a passage of text is to read on a 0-100 scale (higher values indicate greater ease), and the Flesch-Kincaid grade level (Kincaid et al., 1975), which estimates the U.S. school grade required to understand the text. Items demonstrated a standard readability level consistent with plain English (*M* = 62.3, *SD* = 27.5) and were written at a 6^th^- to 7^th^-grade reading level on average (*M* = 6.7 grade, *SD* = 3.63). Although most items were clearly written and accessible, a few were less readable; we revised these to improve clarity and readability. Collectively, Phase 1 resulted in a pool of 67 items that participants interpreted accurately and that experts endorsed as representing the content domain of students mentoring experiences.

### Phase 2: Internal Structure, Convergent, Discriminant, and Criterion-Related Validity

In Phase 2, we collected additional validity evidence of the MERGE as a measure of students’ mentoring experiences. We examined its internal structure using exploratory graph analysis (EGA) and exploratory factor analysis (EFA). We also examined the relationships between MERGE factors and theoretically related constructs (convergent validity), theoretically distinct constructs (discriminant validity), and relevant outcomes (criterion-related validity).

#### Internal Structure Validity

The prior qualitative research on which our items were based indicated that doctoral students described experiencing a range of mentoring experiences, including the presence of supportive behavior (e.g., providing career and psychosocial support), harmful behavior (e.g., yelling, name-calling), and the absence of helpful behaviors (e.g., lack of career guidance, poor availability) (Tuma et al., 2021). As a result, there is some ambiguity regarding how items should be scored. For example, items assessing neglect or insufficient guidance could reflect the low end of a mentoring support continuum (i.e., absence of positive mentoring behavior) rather than a distinct negative mentoring experiences dimension (i.e., presence of harmful mentoring behavior). In this case, responses to negative mentoring items would be expected to cluster with positive mentoring items, reflecting a continuum from high to low support.

The qualitative evidence was unable to establish whether these behaviors reflected opposite ends of a single dimension or distinct dimensions. We therefore treated dimensionality as an empirical question and used hierarchical exploratory graph analysis (hierEGA) to estimate the structure of the item pool without imposing a scoring direction on each item (Jiménez et al., 2025). In addition, a central concern was whether Phase 1 items reflect distinctions in specific mentor behaviors or broader evaluations of the mentoring relationship. We therefore administered five items assessing global mentoring relationship quality (Allen & Eby, 2003) alongside the items generated during Phase 1. The relationship quality items served as a reference point for determining whether the Phase 1 items formed behavioral dimensions distinct from a global appraisal of the relationship. We used the hierEGA solution to inform the number of factors to extract and to specify the target matrix for rotation in the subsequent exploratory factor analyses.

#### Convergent Validity

Convergent validity is demonstrated when responses to different measures of the same or similar constructs are related, reflecting their conceptual overlap (Campbell & Fiske, 1959). Although convergent validity is often tested by examining correlations among multiple distinct operationalizations of a given construct, it can also be exhibited within a measurement model through examining the convergence between different indicators of a given construct (Bagozzi, 1981). Given that the MERGE dimensions are theorized to cohere under a common higher-order mentoring experiences structure, we should expect the dimensions to demonstrate moderate-to-strong associations with each other in theoretically expected directions.

#### Discriminant Validity

Discriminant validity is demonstrated when measures of theoretically distinct constructs are sufficiently different from one another to suggest they capture distinct constructs rather than the same construct (Campbell & Fiske, 1959). We examined discriminant validity in two ways. First, we examined whether the MERGE factors were empirically distinguishable from one another. Because the factors represent various aspects of students’ mentoring experiences, we expected them to be related, but not empirically redundant. To test this, we examined the confidence intervals around the factor correlations to assess the distinctiveness of the factors (Rönkkö & Cho, 2022).

Second, we hypothesized that the MERGE factors would be empirically distinct from students’ personality traits. Given that personality traits are broad and stable dispositions that can influence survey responding (Hibbing et al., 2019), relationship development (Harris & Vazire, 2016), and interpersonal interactions (Mount et al., 1998), scores on the MERGE could reflect students’ personality differences rather than their mentoring experiences. For example, students higher in neuroticism may be more likely to interpret mentors’ benign behavior more negatively; if so, neuroticism would correlate strongly with mentoring experiences. Ratings on personality items also capture individual differences in how individuals respond to surveys, such as “yea-saying” or acquiescing (Danner et al., 2015). We therefore examined associations between the MERGE factors and personality traits in the Five Factor Model: openness, conscientiousness, extraversion, agreeableness, and neuroticism (John & Srivastava, 1999). We hypothesized weak associations, which would provide evidence that the MERGE captures mentoring experiences rather than students’ broad personality traits or response tendencies.

#### Criterion-Related Validity

Criterion-related validity is demonstrated when scores on a measure are associated with theoretically relevant outcomes. We examined whether the MERGE factors were associated with outcomes that we theorized to be related to students’ mentoring experiences using correlation and multiple regression, which we further investigated using dominance analysis.

First, we expected MERGE scores to predict students’ perceptions of support from their graduate program. Perceived organizational support refers to employees’ beliefs that their employer treats them fairly, values their well-being and contributions, and supports their development (Eisenberger et al., 2002). These beliefs develop through employees’ interactions with the organization and its agents (e.g., supervisors and mentors), leading to inferences that the organization values them (or not). Applied to student-faculty mentoring, these processes suggest that mentoring experiences can shape students’ beliefs about the extent to which their program and university care about them. Our prior research provides some evidence that students who experience negative mentoring also feel unsupported or undervalued by their department, program, or university (Tuma et al., 2021). We therefore hypothesized that MERGE factors would predict perceived graduate program support in the expected directions (i.e., positive mentoring experiences would predict higher perceptions of program support, whereas negative mentoring experiences would predict lower support perceptions).

Second, we expected MERGE scores to relate to students’ emotions about their research, including feelings of enthusiasm and enjoyment (i.e., positive affect) as well as frustration and discouragement (i.e., negative affect). Workplace research indicates that employees experience positive emotions when their supervisor provides support and constructive feedback, and negative emotions when they experience inadequate supervision and support (Anttila et al., 2021). Thus, we hypothesized that students’ negative mentoring experiences would be negatively associated with positive affect toward research and positively associated with negative affect toward research. We expected positive mentoring experiences to show the opposite pattern.

#### Phase 2 Methods

##### Participants

Participants were pursuing or had earned a PhD in a life science discipline from a doctoral-granting university in the US and were currently conducting or had conducted doctoral research within the past year. We recruited students directly by email. The recruitment materials described the study as examining the quality of mentorship that graduate students receive and explicitly encouraged students with good, neutral, and poor mentoring experiences to participate. We received 601 responses. We excluded five respondents who did not consent, 16 respondents who completed fewer than 25% of the survey, 15 who failed attention checks, and 35 who met our criteria for careless responding. Following Ward & Meade (2023), we evaluated careless responding across three families of indices: invariability, outlier response patterns, and response consistency. Responses were excluded when responses met criteria across at least two families. The final analytic sample consisted of 530 doctoral students representing 70 institutions across 38 states. Although participants were nested within institutions, intraclass correlation coefficients (ICCs) were uniformly low (<.05), indicating limited shared variance within institutions. In addition, our preliminary attempts to account for clustering resulted in inadmissible solutions because several institutions were represented by only one participant. For these reasons, we did not include institutional clustering in our analytic models.

##### Data Collection

Students who consented to participate were instructed to respond to survey items with their dissertation advisor in mind. The survey asked about the extent to which their advisor acted as a mentor during their graduate research. Participants who were co-advised or had multiple dissertation advisors were asked to select one advisor to report on.

##### Measures

All variables were measured using Likert-type scales (e.g., 1 = “strongly disagree”; 5 = “strongly agree”) and items were coded so that higher values represent greater levels of the construct. Mentoring relationship quality was measured using Allen & Eby’s (2003) five-item scale. Graduate program support was measured using Eisenberger and colleagues’ (1997) eight-item scale of organizational support. Personality traits (agreeableness, conscientiousness, extraversion, openness, neuroticism) were measured using Donnellan and colleagues’ (2006) twenty-item Mini-International Personality Item Pool (mini-IPIP). Participants were asked to think how well the statements described themselves and their personality in general, not as they desired to be in the future. Participants’ positive and negative affect toward research was measured using Watson and colleagues’ (1988) ten-item scale. Participants were asked to think about their graduate research in general and not their mentor as they responded to the items. A complete list of the measures and items are included in the *Supplemental Materials*.

#### Phase 2 Results

##### Exploratory Graph Analyses and Exploratory Factor Analyses

We used exploratory graph analysis (EGA), a network psychometrics approach (Golino et al., 2020), and exploratory factor analysis (EFA) to examine the internal structure of the MERGE. We first used unique variable analysis to identify and remove empirically redundant items (Golino & Christensen, 2024), then we used hierarchical EGA to identify the dimensional structure and inform an iterative series of EFAs (Jiménez et al., 2025). The initial EGA and EFA results indicated that model refinement was needed, so we iteratively combined factors and removed poorly performing items until a stable and interpretable structure was identified. Detailed analytic procedures, item exclusions, and model refinement are described in the *Supplemental Materials*.

The final 37-item measure consists of six factors capturing distinct aspects of doctoral students’ mentoring experiences: *Responsiveness, Career Support, Relationship Strain, Psychosocial Support, Relationship Quality,* and *Negative Mentoring Behaviors*. *Responsiveness* items relate to mentor physical and psychological availability as well as their attention to and engagement with student’s research (e.g., *“My mentor responds when I contact them,” “My mentor gives me their full attention when we meet”). Career Support* items relate to the mentor’s support of the student’s professional development and career advancement (e.g., *“My mentor helps me identify ways to network”*). *Relationship Strain* represents the student’s appraisals of incongruity and tension in the mentoring relationship (e.g., *“My mentor and I have a difficult relationship”*). *Psychosocial Support* reflects the mentor’s encouragement and personal support of the student as well as their attention to the student’s well-being (e.g., *“My mentor empathizes when I am struggling”*). *Relationship Quality* represents student appraisals of the effectiveness and functionality of their mentoring relationship (*“My mentor and I enjoy a high-quality relationship”*). Finally, *Negative Mentoring Behaviors* includes deceitful behavior, abusive supervision, volatility, and boundary-crossing *(“My mentor takes their stress out on me”*). The final network structure is shown in Figure 2, and item loadings are reported in Table 2.

**Figure 2.**
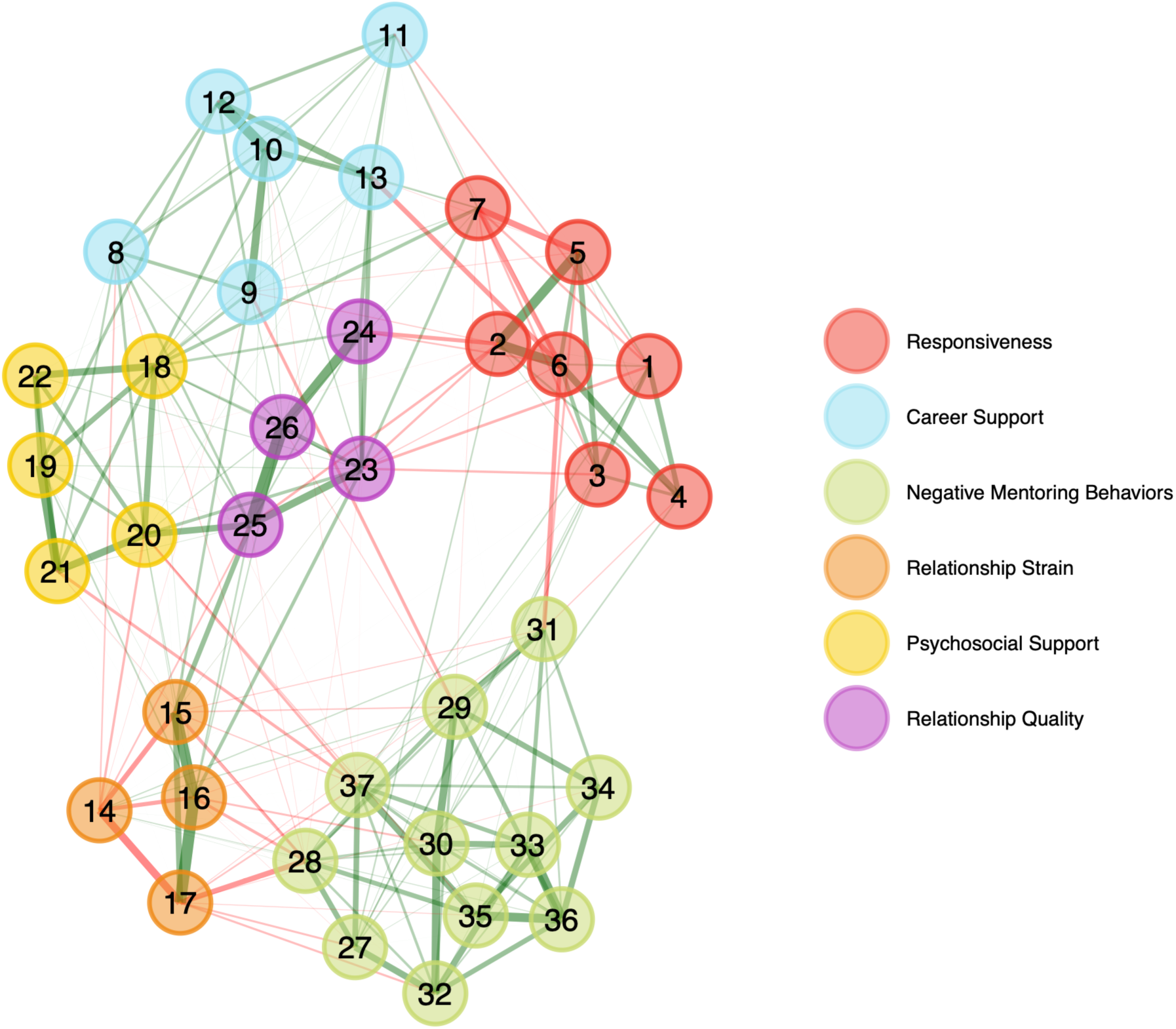
Network Structure of Item Associations. Each node (circle) represents an individual survey item in the final item set (total = 37). Each edge (lines) represents partial correlations between items after controlling for all other items. Distance reflects the strength and uniqueness of the relationships between items. Colors represent empirically derived dimensions (communities) identified by the EGA lower-order algorithm, with items of the same color forming a latent dimension. The resulting clusters represent a stable grouping of items across 500 bootstrapped samples, supporting a six-dimensional lower-order solution.

**Table 2.** Phase 2 Exploratory Factor Analysis Results.

| Item Wording | Responsiveness | Career Support | Relationship Strain | Psychosocial Support | Relationship Quality | Negative Mentoring Behaviors |
| --- | --- | --- | --- | --- | --- | --- |
| My mentor responds when I contact them | **.44** | -.02 | -.04 | .07 | -.00 | -.09 |
| My mentor often forgets about my research progress (r) | **.61** | .01 | .04 | .03 | .21 | -.12 |
| My mentor gives me their full attention when we meet | **.47** | .10 | -.02 | -.13 | .09 | -.20 |
| My mentor's personal demands limit the time they have to mentor me (r) | **.61** | .02 | -.09 | .00 | -.11 | .06 |
| My mentor keeps up with my research progress | **.59** | .10 | -.03 | -.03 | .13 | .02 |
| My mentor is not involved enough in my research (r) | **.81** | .06 | -.06 | .08 | -.05 | .12 |
| My mentor is willing to give me feedback on my research | **.35** | .12 | -.02 | .08 | .17 | -.04 |
| My mentor protects me from others who might cause me professional harm | -.03 | **.36** | -.09 | .16 | .22 | .02 |
| My mentor has little interest in my career advancement (r) | .03 | **.62** | -.17 | -.10 | .04 | -.09 |
| My mentor offers useful advice for achieving my career goals | -.04 | **.95** | -.06 | -.07 | -.06 | -.01 |
| My mentor prioritizes publishing my research | .16 | **.43** | .04 | .01 | .10 | .06 |
| My mentor helps me identify ways to network | -.06 | **.82** | .11 | .11 | -.03 | .02 |
| My mentor helps me prepare for important milestones in my degree | .22 | **.51** | .07 | .09 | .06 | -.07 |
| My mentor and I have incompatible personalities | -.07 | -.07 | **.47** | -.09 | -.07 | .17 |
| My mentor and I do not like each other | .04 | -.13 | **.72** | -.09 | -.08 | .01 |
| My mentor and I have a difficult relationship | -.11 | -.01 | **.70** | -.09 | -.09 | .07 |
| My mentor and I have a tense relationship | -.08 | -.02 | **.67** | -.08 | .02 | .18 |
| My mentor encourages me | -.02 | .20 | -.08 | **.48** | .17 | -.05 |
| My mentor checks in about my wellbeing | .07 | .08 | -.09 | **.69** | -.02 | -.00 |
| My mentor makes me feel accepted | .01 | .08 | -.13 | **.48** | .20 | -.14 |
| My mentor empathizes when I am struggling | -.01 | -.03 | -.06 | **.74** | .08 | -.09 |
| My mentor tells me when they think I have done a good job | .02 | .02 | -.01 | **.67** | .05 | -.04 |
| The mentoring relationship between my mentor and I is very effective | .22 | .13 | -.13 | .15 | **.37** | -.06 |
| I am effectively utilized as a mentee by my mentor | .16 | .10 | .06 | .10 | **.57** | -.04 |
| My mentor and I enjoy a high-quality relationship | .07 | .08 | -.15 | .21 | **.51** | -.05 |
| Both my mentor and I benefit from our mentoring relationship | -.02 | .10 | -.05 | .07 | **.75** | -.04 |
| My mentor says one thing and does another | -.22 | -.04 | .09 | .05 | -.01 | **.57** |
| My mentor is only nice to me when they need me | -.05 | .00 | .35 | -.08 | -.06 | **.43** |
| My mentor threatens me | .04 | .07 | .19 | .13 | -.18 | **.58** |
| My mentor takes their stress out on me | .03 | .12 | .04 | .01 | -.13 | **.78** |
| My mentor micromanages me | .30 | -.13 | .13 | .05 | .03 | **.61** |
| My mentor has an unpredictable personality | -.12 | .08 | -.04 | -.04 | .00 | **.86** |
| My mentor is impatient | .04 | -.03 | -.21 | -.05 | .01 | **.97** |
| My mentor gives me advice on topics that are none of their business | .04 | -.03 | .02 | .14 | -.02 | **.72** |
| My mentor thinks they are better than others | -.03 | -.01 | -.01 | -.21 | .16 | **.76** |
| My mentor is hot tempered | .02 | -.03 | -.07 | .07 | .03 | **.96** |
| My mentor does not take responsibility for their mistakes | -.10 | -.09 | .01 | -.21 | .10 | **.66** |
| Proportion of Variance Explained | 0.09 | 0.10 | 0.09 | 0.09 | 0.08 | 0.19 |
| Omega ($\omega$) Reliability | 0.86 | 0.88 | 0.94 | 0.93 | 0.94 | 0.95 |

*Items marked (r) are reverse scored.*

##### Confirmatory Factor Analysis of Phase 2 Measures

Having established a stable internal structure for the MERGE, we examined associations between its factors and theoretically related constructs. We estimated a combined confirmatory factor analysis (CFA) model including the MERGE factors and the theoretically related constructs, with cross-loadings estimated for the MERGE items. The model demonstrated good fit to the data (χ^2^ (2303) = 3502.49, CFI = 0.96, RMSEA = 0.03, SRMR = 0.04).

##### Evidence of Convergent Validity

Factor correlations supported the hypothesized pattern of associations among the MERGE factors (Table 3). Factors representing positive mentoring experiences (e.g., *Responsiveness, Career Support, Psychosocial Support, Relationship Quality)* were positively associated with one another, with latent correlations ranging from .55 to .76. For example, *Career Support* was strongly associated with *Psychosocial Support* (φ = 0.76) and *Relationship Quality* (φ = 0.75). *Relationship Strain* and *Negative Mentoring Behaviors* were also strongly positively associated (φ = 0.79), consistent with their shared representation of negative mentoring experiences. In contrast, factors representing negative mentoring experiences were negatively associated with factors representing positive mentoring experiences, with latent correlations ranging from -.39 to -.66. All associations were in the expected directions, providing evidence of convergent validity.

**Table 3.**
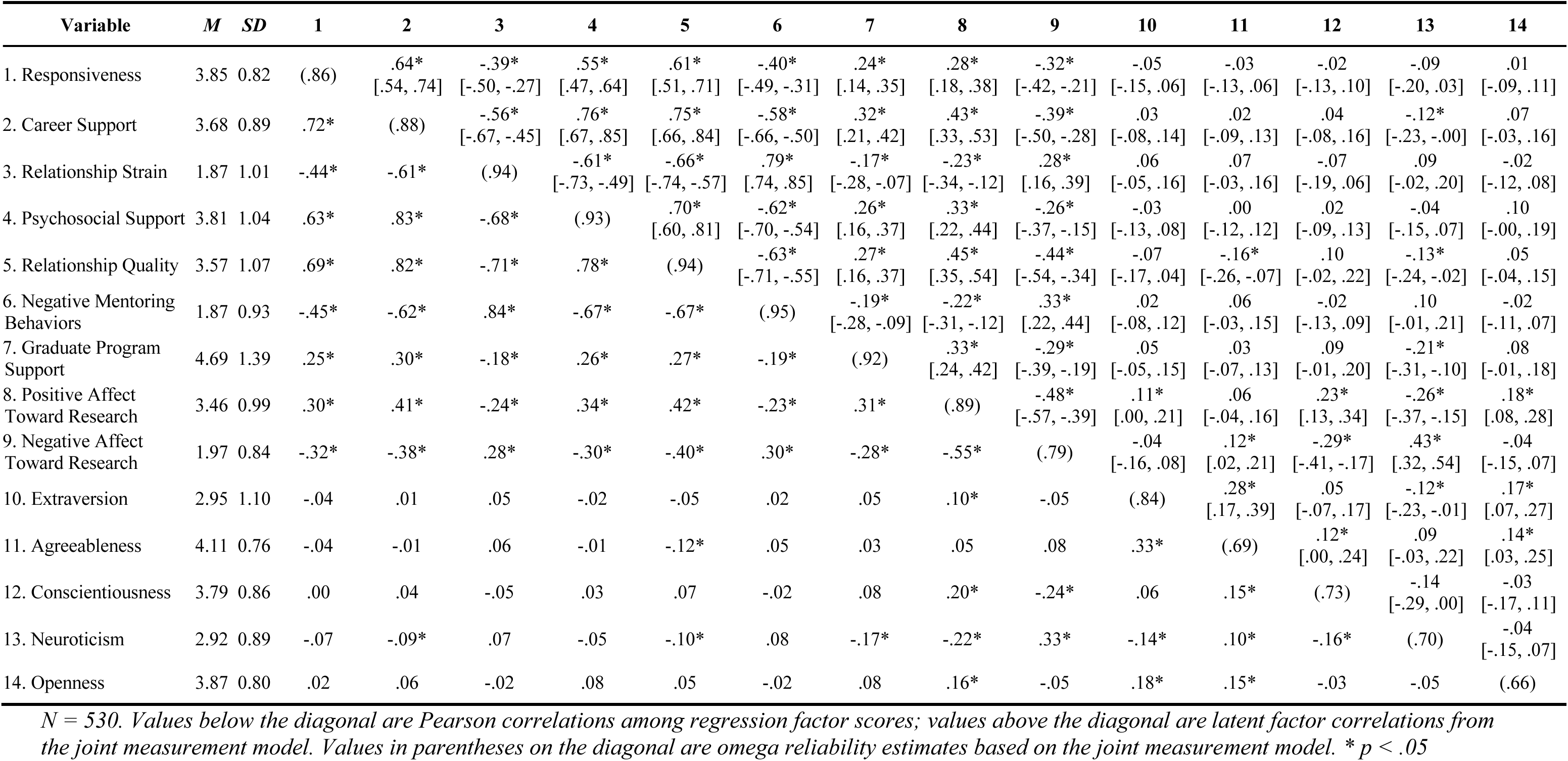
Phase 2 Variable Correlations.

##### Evidence of Discriminant Validity

Following Rönkkö & Cho (2022), we evaluated discriminant validity by examining the confidence intervals around the latent factor correlations (Table 3). Correlations among the six MERGE factors ranged from |φ| = .39 to .79, and the confidence intervals did not indicate problematic overlap between factors. These results indicate that the six MERGE factors are meaningfully related to one another but empirically distinguishable dimensions of doctoral students’ mentoring experiences.

Consistent with expectations, the MERGE factors showed very weak associations with personality traits (i.e., latent correlations ranging from .00 to .16), indicating that reports of mentoring experiences captured by the MERGE are largely independent of personality traits and individual differences in response tendencies.

##### Evidence of Criterion-Related Validity

All MERGE factors were significantly associated with graduate program support (φ = -.17 to .32), positive affect toward research (φ = -.23 to .45), and negative affect toward research (φ = -.44 to .33) in the expected directions, providing evidence of criterion-related validity. We also used dominance analyses to examine the relative contribution of the six MERGE factors to each criterion variable while accounting for overlap among factors (Table 4). Dominance analysis is a variance decomposition technique that estimates the relative importance of correlated predictors by evaluating their contribution to explained variance (*R^2^*) across all subsets of predictors in regression models (Azen & Budescu, 2003). Detailed analytic procedures are provided in the *Supplemental Materials*. The six MERGE factors jointly explained 9% of the variance in graduate program support, 20% of the variance in positive affect toward research, and 18% of the variance in negative affect toward research.

**Table 4.** Phase 2 Dominance Analysis.

| Predictor |  |  |  | Outcome |  |  |  |  |  |
| --- | --- | --- | --- | --- | --- | --- | --- | --- | --- |
| MERGE Factor | Graduate Program Support |  |  | Positive Affect Toward Research |  |  | Negative Affect Toward Research |  |  |
| | $\beta$ | Dominance | sr<sup>2</sup> | $\beta$ | Dominance | sr<sup>2</sup> | $\beta$ | Dominance | sr<sup>2</sup> |
| Responsiveness | 0.05 | 0.02 | 0.00 | -0.05 | 0.02 | 0.00 | -0.04 | 0.02 | 0.00 |
| Career Support | 0.21* | **0.03** | 0.01 | 0.28* | 0.06 | 0.02 | -0.20* | 0.04 | 0.01 |
| Relationship Strain | 0.04 | 0.01 | 0.00 | 0.06 | 0.01 | 0.00 | -0.09 | 0.01 | 0.00 |
| Psychosocial Support | 0.04 | 0.02 | 0.00 | -0.03 | 0.03 | 0.00 | 0.17* | 0.02 | 0.01 |
| Relationship Quality | 0.05 | 0.02 | 0.00 | 0.35* | **0.07** | 0.03 | -0.32* | **0.06** | 0.02 |
| Negative Mentoring Behaviors | -0.00 | 0.01 | 0.00 | 0.09 | 0.01 | 0.00 | 0.14 | 0.02 | 0.01 |
| **R<sup>2</sup>** | **0.09** |  |  | **0.20** |  |  | **0.18** |  |  |

*N = 530. Beta ( $\beta$ ) is the standardized OLS coefficient from a simultaneous regression of the outcome on all 6 MERGE factor scores; \* $p < .05$ . Dominance is the general dominance weight (each predictor's marginal contribution to $R^2$ across every possible subset of the other predictors, summing to the model $R^2$ ). sr<sup>2</sup> is the semi-partial $R^2$ (the drop in model $R^2$ if that predictor alone were removed and the other five retained). Bolded Dominance values indicate the highest-Dominance predictor for the outcome.*

For graduate program support, *Career Support* dominated the other MERGE factors in relative importance (dominance = .03), accounting for 3% of the variance in graduate program support or one-third of the total variance in graduate program support explained by the MERGE. *Career Support* also accounted for the largest unique share of variance in graduate program support (sr^2^ =.01). Career support was significantly and positively associated with graduate program support in the simultaneous regression model (β = .21). The remaining MERGE factors had smaller dominance weights (.01 - .02) and did not independently predict graduate program support when career support was included in the model. For positive affect toward research, *Relationship Quality* exhibited general dominance over the other MERGE factors (dominance = .07), followed by *Career Support* (dominance = .06). *Relationship Quality* also accounted for the largest unique share of variance (sr² = .03), followed by *Career Support* (sr² = .02). Both factors were positively associated with positive affect toward research when the other MERGE factors were included in the model (β = .35 and .28, respectively). For negative affect toward research, *Relationship Quality* again exhibited general dominance over the other MERGE factors (dominance = .06), followed by *Career Support* (dominance = .04). *Relationship Quality* accounted for the largest unique share of variance (sr² = .02) and was negatively associated with negative affect toward research (β = −.32). *Career Support* was similarly negatively associated with negative affect toward research (β = −.20) and accounted for .01 of unique variance. *Psychosocial Support* was positively associated with negative affect toward research (β = .17) when controlling for the other MERGE factors but had a smaller relative contribution (dominance = .02; sr² = .01).

In sum, the pattern of associations between the MERGE factors and criterion variables was consistent with our hypotheses, providing evidence for the criterion-related validity of the MERGE. The dominance analyses further showed that the MERGE factors did not contribute equally to these outcomes: *Career Support* was the most important factor for predicting graduate program support, whereas *Relationship Quality* was the most important factor for predicting both positive and negative affect toward research. These results indicate that the MERGE factors are meaningfully related to theoretically relevant variables while also revealing differences in the relative importance of specific mentoring experiences for different student outcomes.

### Phase 3: Replication of Factor Structure, Criterion-Related Validity, and Incremental Validity

In Phase 3, we assessed the replicability of the Phase 2 MERGE factor structure using a new, independent sample. This sample included doctoral students outside of the life sciences to enable examination of the generalizability of the factor structure beyond the Phase 2 sample. We further evaluated the criterion-related validity of the MERGE by examining associations between MERGE scores and student reports of their career and well-being-related outcomes. Finally, we examined whether the MERGE explained variance in student outcomes above and beyond an existing measure of mentor competency (i.e., incremental validity).

#### Factor Structure Replication

Replicating the factor structure in an independent sample provides evidence that the observed dimensionality is not unique to the original sample and increases confidence in the stability of the measure (Floyd & Widaman, 1995).

#### Criterion-Related Validity

We further examined associations between the MERGE factors and additional theoretically relevant career and well-being-related outcomes relevant to doctoral students, including research self-efficacy, burnout, work-family conflict, anxiety, and career intentions. The rationale and hypothesis for each criterion variable are provided in the *Supplemental Materials*.

#### Incremental Validity

If the MERGE captures aspects of mentoring that are not adequately represented by existing measures, it should explain additional variance in relevant outcomes. To test this, we examined the incremental validity of the MERGE relative to the *Mentoring Competency Assessment (MCA)* (Fleming et al., 2013), a commonly used measure of mentoring. The MCA was developed to assess mentors’ competencies across six domains of effective mentoring addressed in the professional development program, *Entering Mentoring* (Handelsman et al., 2011). The MCA has been used to query mentors about their own competencies and query students about their mentors’ competencies but is not designed to assess dysfunctional or harmful mentoring behaviors. Predicting student outcomes with both the MERGE and the MCA using the same sample provides a test of whether assessing a broader range of mentoring experiences, including positive and negative experiences, adds value when predicting student outcomes.

#### Phase 3 Methods

##### Participants

Participants were pursuing or had earned a PhD in a natural science (e.g., biology, chemistry, computer science, geology) or engineering discipline at a US doctoral-granting university and were currently conducting or had conducted doctoral research for at least one year within the past year. As in Phase 2, we recruited students directly and encouraged individuals with good, neutral, or poor mentoring experiences to participate. We received 861 responses. We excluded eight respondents who did not consent, 48 who completed less than 25% of the survey, and 15 who failed attention checks. We also screened for careless responding using the same procedures as Phase 2, resulting in the exclusion of 41 additional responses. The final analytic sample consisted of 749 doctoral students representing 73 public and private institutions across 39 states.

##### Data Collection

Participants were instructed to think about their dissertation advisor and respond to the survey items with this individual in mind. As in Phase 2, participants who were co-advised or had multiple dissertation advisors were asked to select one advisor to report on.

##### Measures

All variables were measured using five-point Likert scales (i.e., 1 = “strongly disagree”; 5 = “strongly agree”) and items were coded so that higher values represent greater levels of the construct unless otherwise noted. Student perceptions of their mentor’s competencies were measured using Hyun and colleagues’ (2022) 21-item Revised Mentoring Competency Assessment (i.e., MCA-21). Participants rated their mentor’s skill in each behavior (1 = not at all skilled to 7 = extremely skilled). Career intentions were measured using Estrada and colleagues’ (2019) four-item scale. Participants indicated the likelihood of pursuing each career path (1 = not at all likely to 5 = absolutely likely). Research self-efficacy was measured using Kardash’s (2000) 13-item scale^1^. Participants rated their confidence in performing research tasks (1 = not at all confident to 5 = very confident). Anxiety was measured using Spitzer and colleagues’ (2006) seven-item Generalized Anxiety Disorder-7 scale, a widely used screening measure of anxiety symptoms in clinical settings. Participants indicated how often (1 = not at all to 4 = nearly every day) they had been bothered by a series of anxiety symptoms over the past two weeks. Burnout was measured using Salmela-Aro and colleagues’ (2009) nine-item School Burnout Inventory. Participants rated their agreement (1 = completely disagree to 6 = completely agree) with a series of items assessing multiple facets of burnout, including cynicism, exhaustion, and sense of inadequacy. Work-family conflict was measured using Netemeyer and colleagues’ (1996) five-item scale. A complete list of the measures and items are included in the *Supplemental Materials*.

#### Phase 3 Results

##### Replication of the MERGE Factor Structure

To evaluate the replicability of the MERGE factor structure, we estimated the six-factor model identified in Phase 2 with the Phase 3 sample comprised of natural science and engineering doctoral students. The resulting solution reproduced the pattern observed^2^ in Phase 2 and demonstrated excellent fit, χ² (459) = 873.56, *p*

< 0.001, CFI = 0.98, RMSEA = 0.04, SRMR = .02. These results indicate that the internal structure of the MERGE is replicable and consistent for natural science and engineering doctoral students.

##### Criterion-Related Validity

As hypothesized, all six MERGE factors were significantly associated with career- and wellbeing-related outcomes (Table 5). Specifically, all MERGE factors were significantly associated with work-family conflict (φ = -.38 to .37), anxiety (φ = -.32 to .35), burnout (φ = -.58 to .50), and career intentions (φ = -.29 to .34) in the expected directions. All factors were also significantly associated with research self-efficacy (φ = -.10 to .30), except *Relationship Strain* (φ = -.06, ns). These results provide additional evidence of criterion validity of the MERGE.

**Table 5.** Phase 3 Variable Correlations.

| Variable | *M* | *SD* | 1 | 2 | 3 | 4 | 5 | 6 | 7 | 8 | 9 | 10 | 11 | 12 |
| --- | --- | --- | --- | --- | --- | --- | --- | --- | --- | --- | --- | --- | --- | --- |
| 1. Responsiveness | 3.86 | 0.87 | (.88) | .50*<br>[.42, .59] | -.34*<br>[-.41, -.26] | .48*<br>[.40, .55] | .67*<br>[.60, .73] | -.50*<br>[-.57, -.43] | -.21*<br>[-.30, -.13] | -.21*<br>[-.29, -.12] | .29*<br>[.20, .38] | -.43*<br>[-.51, -.35] | .18*<br>[.10, .27] | .70*<br>[.65, .75] |
| 2. Career Support | 3.68 | 0.95 | .78* | (.89) | -.49*<br>[-.56, -.41] | .51*<br>[.37, .65] | .58*<br>[.48, .68] | -.47*<br>[-.55, -.39] | -.20*<br>[-.30, -.11] | -.13*<br>[-.23, -.04] | .33*<br>[.22, .43] | -.33*<br>[-.43, -.24] | .21*<br>[.10, .31] | .72*<br>[.64, .81] |
| 3. Relationship Strain | 1.97 | 1.05 | -.73* | -.84* | (.94) | -.49*<br>[-.60, -.37] | -.56*<br>[-.66, -.46] | .71*<br>[.63, .79] | .26*<br>[.17, .34] | .25*<br>[.16, .33] | -.23*<br>[-.33, -.14] | .37*<br>[.28, .46] | -.06<br>[-.16, .04] | -.59*<br>[-.68, -.49] |
| 4. Psychosocial Support | 3.76 | 1.05 | .77* | .92* | -.93* | (.92) | .70*<br>[.63, .76] | -.67*<br>[-.75, -.59] | -.38*<br>[-.48, -.29] | -.29*<br>[-.38, -.20] | .30*<br>[.21, .40] | -.50*<br>[-.58, -.42] | .15*<br>[.06, .24] | .80*<br>[.75, .85] |
| 5. Relationship Quality | 3.46 | 1.16 | .81* | .89* | -.90* | .93* | (.95) | -.68*<br>[-.74, -.63] | -.28*<br>[-.36, -.20] | -.32*<br>[-.39, -.24] | .34*<br>[.26, .42] | -.58*<br>[-.65, -.52] | .30*<br>[.22, .38] | .86*<br>[.83, .89] |
| 6. Negative Mentoring Behaviors | 1.99 | 0.98 | -.69* | -.79* | .89* | -.87* | -.83* | (.94) | .37*<br>[.30, .44] | .35*<br>[.28, .42] | -.29*<br>[-.37, -.21] | .50*<br>[.44, .56] | -.10*<br>[-.18, -.02] | -.77*<br>[-.81, -.74] |
| 7. Work-Family Conflict | 3.12 | 1.11 | -.26* | -.32* | .34* | -.35* | -.31* | .36* | (.94) | .49*<br>[.43, .56] | -.15*<br>[-.24, -.06] | .56*<br>[.50, .62] | -.04<br>[-.12, .05] | -.35*<br>[-.42, -.28] |
| 8. Anxiety | 1.20 | 0.91 | -.25* | -.27* | .32* | -.31* | -.32* | .34* | .47* | (.93) | -.15*<br>[-.24, -.06] | .66*<br>[.61, .72] | -.15*<br>[-.23, -.07] | -.30*<br>[-.38, -.23] |
| 9. Career Intentions | 3.34 | 1.00 | .29* | .34* | -.29* | .32* | .32* | -.28* | -.13* | -.13* | (.75) | -.43*<br>[-.50, -.36] | .28*<br>[.19, .36] | .39*<br>[.31, .47] |
| 10. Burnout | 3.55 | 1.18 | -.47* | -.51* | .52* | -.53* | -.56* | .50* | .51* | .61* | -.37* | (.90) | -.29*<br>[-.37, -.21] | -.57*<br>[-.62, -.51] |
| 11. Research Self-Efficacy | 4.07 | 0.57 | .19* | .22* | -.15* | .18* | .24* | -.12* | -.03 | -.14* | .23* | -.26* | (.91) | .27*<br>[.19, .35] |
| 12. Mentor Competency Assessment (MCA-21) | 4.57 | 1.64 | .78* | .89* | -.82* | .89* | .89* | -.80* | -.33* | -.29* | .34* | -.53* | .26* | (.98) |

*N* = 747-749 per variable due to a small number of cases missing. Values below the diagonal are Pearson correlations among regression factor scores; values above the diagonal are latent factor correlations from the joint measurement model. Values in parentheses on the diagonal are omega reliability estimates for each composite. \**p* < .05

To further investigate the unique effects of each MERGE factor, we conducted dominance analysis^3^ (Table 6) for each outcome, including assessment of independent associations for each factor (i.e., β). We examined two career-related outcomes: research self-efficacy and career intentions. *Relationship Quality* dominated the other MERGE factors in relative importance (dominance = .04) when predicting self-efficacy, followed by *Career Support* (dominance =.02). *Relationship Quality* was positively associated with self-efficacy (β = .56) and accounted for the largest unique share of variance (sr² = .03). *Career Support* was also positively associated with self-efficacy (β = .20) and accounted for .01 of unique variance. *Negative Mentoring Behaviors* was also positively associated with self-efficacy (β =.17) but demonstrated less unique predictive power (dominance = .01; sr² = .01). For career intentions, *Career Support* dominated the other MERGE factors in relative importance (dominance = .03). *Career Support* was positively associated with career intentions (β = .28) and accounted for the largest unique share of variance (sr² = .01). The remaining MERGE factors had small conditional associations with *Career Intentions* (βs = -.04 to .05) and accounted for no unique variance when controlling for the other MERGE factors (sr² = .00).

**Table 6.** Phase 3 Dominance Analysis.

| Predictor |  |  |  | Outcome |  |  |  |  |  |  |  |  |  |  |  |
| --- | --- | --- | --- | --- | --- | --- | --- | --- | --- | --- | --- | --- | --- | --- | --- |
| MERGE Factor | Work-Family Conflict |  |  | Anxiety |  |  | Career Intentions |  |  | Burnout |  |  | Self-Efficacy |  |  |
| | $\beta$ | Dominance | sr<sup>2</sup> | $\beta$ | Dominance | sr<sup>2</sup> | $\beta$ | Dominance | sr<sup>2</sup> | $\beta$ | Dominance | sr<sup>2</sup> | $\beta$ | Dominance | sr<sup>2</sup> |
| Responsiveness | 0.01 | 0.01 | 0.00 | 0.02 | 0.01 | 0.00 | 0.04 | 0.02 | 0.00 | -0.06 | 0.04 | 0.00 | -0.02 | 0.01 | 0.00 |
| Career Support | -0.00 | 0.02 | 0.00 | 0.15 | 0.01 | 0.00 | 0.28* | **0.03** | 0.01 | 0.00 | 0.05 | 0.00 | 0.20* | 0.02 | 0.01 |
| Relationship Strain | 0.00 | 0.02 | 0.00 | 0.02 | 0.02 | 0.00 | 0.01 | 0.01 | 0.00 | 0.01 | 0.05 | 0.00 | 0.12 | 0.01 | 0.00 |
| Psychosocial Support | -0.32* | 0.03 | 0.01 | -0.03 | 0.02 | 0.00 | -0.04 | 0.02 | 0.00 | 0.00 | 0.05 | 0.00 | -0.25 | 0.01 | 0.00 |
| Relationship Quality | 0.17 | 0.02 | 0.00 | -0.22* | 0.02 | 0.00 | 0.05 | 0.02 | 0.00 | -0.42* | **0.08** | 0.02 | 0.56* | **0.04** | 0.03 |
| Negative Mentoring Behaviors | 0.22* | **0.03** | 0.01 | 0.24* | **0.03** | 0.01 | -0.03 | 0.01 | 0.00 | 0.10 | 0.05 | 0.00 | 0.17* | 0.01 | 0.01 |
| **R<sup>2</sup>** | **0.14** |  |  | **0.12** |  |  | **0.12** |  |  | **0.31** |  |  | **0.09** |  |  |

*N* = 747-749 (varies slightly by outcome due to a small number of cases missing). Beta ( $\beta$ ) is the standardized OLS coefficient from a simultaneous regression of the outcome on all 6 MERGE factor scores; \* $p < .05$ . Dominance is the general dominance weight (each predictor's marginal contribution to $R^2$ across every possible subset of the other predictors, summing to the model $R^2$ ). $sr^2$ is the semi-partial $R^2$ (the drop in model $R^2$ if that predictor alone were removed while retaining the other five). Bolded Dominance values indicate the highest-Dominance predictor for the outcome.

We used MERGE factors to predict three well-being outcomes: anxiety, burnout, and work-family conflict. For anxiety, *Negative Mentoring Behaviors* exhibited dominance over the other MERGE factors (dominance = .03), followed by *Relationship Quality, Relationship Strain,* and *Psychosocial Support* (dominance =.02). *Negative Mentoring Behaviors* had a positive independent association with anxiety (β = .24) and accounted for the largest unique share of variance (sr² = .01). *Relationship Quality* was negatively associated with anxiety (β = -.22) but accounted for little unique variance (sr² = .00). The remaining factors had small associations (βs = -.03 to .15) and little unique contribution. For burnout, *Relationship Quality* dominated the other MERGE factors in relative importance (dominance = .08). *Relationship Quality* was negatively associated with burnout (β = -.42) and accounted for the largest unique share of variance (sr² = .02). The remaining MERGE factors demonstrated smaller dominance weights (.04 - .05) for burnout and accounted for no unique variance (sr² = .00). *Psychosocial Support* and *Negative Mentoring Behaviors* dominated the other MERGE factors in relative importance

(dominance = .03 for both) for predicting work-family conflict, accounting for the largest share of explained variance. *Psychosocial Support* had a negative association with work-family conflict (β = -.32), whereas *Negative Mentoring Behaviors* had a positive association (β = .22). Both accounted for a small amount of unique variance (sr² = .01). The remaining MERGE factors demonstrated little relative or unique contribution to work-family conflict.

Considered together, the findings provide evidence of criterion-related validity while illustrating variation in the associations between different dimensions of the MERGE and student outcomes. Although the bivariate associations showed that nearly all MERGE factors were significantly associated with the criterion variables, the dominance analyses indicated differences in their relative and unique contributions across outcomes. *Relationship Quality* demonstrated the greatest relative importance for predicting self-efficacy and burnout, *Career Support* demonstrated the greatest relative importance for predicting career intentions, and *Negative Mentoring Behaviors* demonstrated the greatest relative importance for predicting anxiety. For work-family conflict, *Psychosocial Support* and *Negative Mentoring Behaviors* demonstrated the greatest relative importance, with the former negatively and the latter uniquely positively associated with work-family conflict. These results highlight the value of distinguishing among dimensions of mentoring when examining their associations with career and well-being outcomes and provide further support for the multidimensional structure of the MERGE.

##### Incremental Validity

We used hierarchical multiple regression to determine whether the MERGE explained additional variance in outcomes beyond the MCA-21. We represented the MCA-21 as a single second-order total score, as its six lower-order factors were highly correlated (latent correlations = .79 to .95). Results modeling the MCA-21 at the individual factor level are reported in the *Supplemental Materials.* For each outcome, we compared models containing the MCA-21 alone, the six MERGE factors alone, and both measures together.

The MERGE demonstrated incremental validity beyond the MCA-21 for four of the five outcomes examined (Table 7). Specifically, the MERGE explained additional variance in work-family conflict, anxiety, burnout, and self-efficacy beyond the MCA-21, with increases in adjusted *R²* ranging from .03 to .04. In contrast, the MCA-21 explained significant additional variance over the MERGE for only one outcome (based on the 95% C.I.). Specifically, the MCA-21 explained additional variance in self-efficacy beyond the MERGE (Δadj. *R^2^* = .02), but this increment was smaller than that uniquely predicted by the MERGE (Δadj. *R^2^* = .04). Neither measure demonstrated clear incremental validity over the other for career intentions. Detailed model comparisons, confidence intervals, and interpretation are provided in the *Supplemental Materials*. The pattern of results indicated that the MERGE captured outcome variance not accounted for by the MCA-21.

**Table 7.** Phase 3 Incremental Validity: MERGE vs. MCA-21.

| Outcome | *N* | *R*<sup>2</sup><br>MCA-only | *R*<sup>2</sup><br>MERGE-only | *R*<sup>2</sup><br>Combined | MERGE incremental variance over MCA-21 |  |  |  | MCA-21 incremental variance over MERGE |  |  |  |
| --- | --- | --- | --- | --- | --- | --- | --- | --- | --- | --- | --- | --- |
| | | | | | $\Delta$ adj.*R*<sup>2</sup> | *F* | *p* | 95% *CI* | $\Delta$ adj.*R*<sup>2</sup> | *F* | *p* | 95% *CI* |
| Work-Family Conflict | 748 | 0.11 | 0.13 | 0.13 | **0.03** | F(6, 740) = 4.81 | < .001 | [0.01, 0.07] | 0.00 | F(1, 740) = 2.37 | 0.124 | [-0.00, 0.01] |
| Anxiety | 747 | 0.08 | 0.12 | 0.11 | **0.03** | F(6, 739) = 5.44 | < .001 | [0.01, 0.07] | -0.00 | F(1, 739) = 0.33 | 0.566 | [-0.00, 0.01] |
| Career Intentions | 748 | 0.11 | 0.11 | 0.11 | -0.00 | F(6, 740) = 1.00 | 0.425 | [-0.00, 0.03] | 0.00 | F(1, 740) = 5.00 | 0.026 | [-0.00, 0.02] |
| Burnout | 749 | 0.28 | 0.31 | 0.31 | **0.03** | F(6, 741) = 6.66 | < .001 | [0.02, 0.07] | 0.00 | F(1, 741) = 6.21 | 0.013 | [-0.00, 0.02] |
| Self-Efficacy | 749 | 0.06 | 0.08 | 0.10 | **0.04** | F(6, 741) = 6.07 | < .001 | [0.02, 0.08] | **0.02** | F(1, 741) = 14.66 | < .001 | [0.00, 0.04] |

*R*<sup>2</sup> values are adjusted *R*<sup>2</sup>. Δadj.*R*<sup>2</sup> is the increment in adjusted *R*<sup>2</sup> from the restricted model (MCA-only for the MERGE-increment columns; MERGE-only for the MCA-increment columns) to the combined model; bolded values have a 1000-rep bootstrap 95% *CI* excluding 0. *F* is the nested-model *F*-test on the *R*<sup>2</sup> increment.

## DISCUSSION

The objectives of this research were to (a) advance the conceptualization of mentoring experiences by capturing the range of effective and negative experiences doctoral students encounter in their mentoring relationships with faculty, (b) develop a psychometrically sound measure that distinguishes among these dimensions, and (c) evaluate the validity of the resulting measure. These efforts resulted in the development of the 37-item MERGE scale and evidence of its validity as a measure of doctoral students’ mentoring experiences. Across three phases and a combined sample of more than 1,300 doctoral students, we found evidence supporting the MERGE’s validity across domains, including content, response process, readability, internal structure, relations with other variables, criterion-related, and incremental validity. Our results advance conceptualization and measurement of the range of mentoring experienced by doctoral students that can be used to advance theory, research, and practice.

The MERGE captures six unique dimensions of doctoral students’ mentoring experiences, from supportive to negative: *Responsiveness, Career Support, Relationship Strain, Psychosocial Support, Relationship Quality, and Negative Mentoring Behaviors.* This conceptualization is broader than existing measures and recognizes that mentoring relationships can involve multiple types of experiences that are not uniformly positive or negative. For example, a student may receive sufficient career support while also experiencing relationship strain. Measures that focus primarily on positive or supportive aspects of mentorship are likely to provide an incomplete account of students’ experiences, and as a result are likely to be less predictive of students’ outcomes. The MERGE provides a way to examine this broader range of experiences, including their antecedents, correlates, and consequences.

Our findings provide evidence that negative mentoring experiences are conceptually and empirically distinct from supportive mentoring experiences. Both *Negative Mentoring Behaviors* and *Relationship Strain* factored separately from supportive mentoring experiences, consistent with prior conceptualizations of negative mentoring as involving relational processes that extend beyond the absence of positive mentoring functions (Eby et al, 2000; 2004). *Negative Mentoring Behaviors* and *Relationship Strain* were also strongly correlated (φ = 0.79 and 0.71 in Phases 2 and 3, respectively) but not redundant. This correlation is not surprising given that relationship strain may arise because of problematic mentor behaviors but may also reflect students’ broader appraisals of the relationship. Our results provide evidence that negative mentoring experiences include distinguishable appraisals of mentor behaviors and mentoring relationship difficulties. Although negative aspects of mentoring were distinct, some aspects traditionally characterized as negative mentoring (e.g., neglect) may reflect the low end of responsiveness - a construct that is well established in relationship science but largely absent from conceptualizations of mentoring (Feeney & Collins, 2015; Reis & Gable, 2015).

The MERGE has multiple advantages over a widely used measure (i.e., MCA-21) for examining research and practice on mentoring. The MCA-21 factors were highly intercorrelated, consistent with prior work showing substantial overlap among factors of both the full MCA and the MCA-21 (Fleming et al., 2013; Zhong & Maccalla, 2024). Such overlap makes it difficult to distinguish how individual mentoring competencies uniquely contribute to student outcomes. In contrast, the MERGE factors are empirically distinguishable and show unique predictive patterns for student outcomes, offering more targeted insight into which aspects of students’ mentoring experiences relate to each outcome. Furthermore, the MERGE captures both positive and negative mentoring experiences within a single measure, offering a comprehensive “toolkit” for examining the range of doctoral student mentoring experiences.

Our work highlights what can be learned by distinguishing among the range of mentoring experienced by students and how these experiences relate to various career and well-being outcomes. The dominance analyses revealed that the six MERGE factors did not contribute equally to outcomes, and no single factor consistently demonstrated the greatest relative importance across outcomes. For example, *Relationship Quality* contributed the most to research self-efficacy and burnout, while *Career Support* contributed most to career intentions. *Negative Mentoring Behaviors* uniquely contributed the most to anxiety and was among the largest contributors to work-family conflict. Collectively, these results show that it is both important and feasible to distinguish among the different components of mentorship rather than treating students’ mentoring experiences as an undifferentiated construct. The MERGE is a tool that can help move the field beyond asking whether mentoring is broadly associated with student outcomes toward identifying *which* aspects of mentoring are most explanatory or predictive. Distinguishing among these aspects of mentoring can enable a more precise accounting of how aspects of mentoring relate to doctoral students’ experiences and reveal features of mentoring that warrant greater attention.

## LIMITATIONS AND FUTURE RESEARCH

The current study has several limitations that also highlight opportunities for future research. The data were collected using a cross-sectional design, which precludes causal inference and prevents establishment of a temporal order. Reverse causality and bidirectional effects are possible. For example, students who struggle to balance the demands of doctoral training may influence mentors’ support and, in turn, students’ perceptions of their mentoring experiences. Conversely, differences in mentoring support may shape students’ perceptions of their ability to manage those demands. Longitudinal research is needed to examine these potentially reciprocal processes and understand how mentoring experiences change over time.

Although this study provides evidence supporting the validity of the MERGE, additional evidence is needed before using the measure for high-stakes decision making. For example, practitioners may be keen to use the MERGE to reward highly effective mentoring or correct problematic mentoring. However, we did not establish meaningful score interpretations or examine what levels of mentoring experiences warrant recognition or intervention (e.g., cut scores). The consequences of using the MERGE also need to be considered. For example, using MERGE scores to evaluate individual mentors could influence their motivation to mentor, particularly if scores are related to formal evaluation, rewards, or sanctions. Students may experience retaliation if their MERGE responses are used in evaluative decisions involving their mentors. Faculty may also suffer unintended consequences if students respond to the MERGE with an intent to harm them rather than to honestly report their experiences. Future research should examine appropriate use of MERGE scores and explore the potential for unintended negative consequences.

Although prior research suggests that negative mentoring experiences are multidimensional (Tuma et al., 2021), the MERGE captures specific experiences primarily within a broad *Negative Mentoring Behaviors* factor. Our initial EGA results provide some evidence of finer-grained subdimensions within the negative mentoring items. However, these distinctions did not persist in the EFA because several items were similarly worded and because potential subdimensions were represented by too few items. The current *Negative Mentoring Behaviors* factor may therefore represent several forms of negative mentoring experiences that could not be reliably differentiated here. Future research could explore a more fine-grained assessment of the lower-order factors reflected by negative mentoring experiences.

All data collected in this study were self-reported by a single source (i.e., the mentee). Although perceptions of mentoring experiences are predictive of mentee outcomes and therefore important to measure, relying on one source may introduce common method variance, recall bias, and attributional biases. Such biases are important to consider given that mentee and mentor reports of the same experiences are not highly correlated (Fagenson-Eland et al., 2005; Welsh & Diehn, 2018). Future research could consider incorporating behavioral observations to assess mentoring experiences.

## ACKNOWLEDGMENTS

We thank Melissa Aikens, Peggy Brickman, Angela Byars-Winston, Lisa Corwin, Lillian Eby, Kimberly Griffin, Paul Hernandez, Becca Price, and Christiane Spitzmueller for their feedback. This work was supported by funding from a National Science Foundation (NSF), Graduate Research Fellowship for TTT (Division of Graduate Education [DGE] grant no. 1842396), NSF DGE grant no. 2328692, and the Georgia Athletic Association Professorship for Innovative Science Education. Any opinions, findings, conclusions, or recommendations expressed in this material are those of the authors and do not necessarily reflect the views of the NSF.

## Footnotes

1 The original version of this scale includes 14 items. One item (“*identify a specific research question for investigation based on the research in your field*”) was inadvertently omitted from the survey during data collection.

2 36 of the 37 items loaded most strongly on their intended factor. The exception was MERGE1 (“*My mentor is willing to give me feedback on my research”*) which loaded on both *Career Support* (.43) and *Responsiveness* (.31). The item’s content spans aspects of both *Career Support* and *Responsiveness,* which may account for its loading pattern.

3 β represents the standardized regression coefficient for each MERGE factor when all six factors were included simultaneously; therefore, β reflects the individual factor’s association with the outcome after accounting for overlap with the other MERGE factors.

## REFERENCES

1. Allen, T. D., & Eby, L. T. (2003). Relationship Effectiveness for Mentors: Factors Associated with Learning and Quality. Journal of Management, 29(4), 469–486.

2. Allen, T. D., & Eby, L. T. (Eds.). (2007). The Blackwell Handbook of Mentoring (p. g9781405133739toc). Blackwell Publishing Ltd.

3. Allen, T. D., Eby, L. T., Poteet, M. L., Lentz, E., & Lima, L. (2004). Career Benefits Associated With Mentoring for Proteges: A Meta-Analysis. Journal of Applied Psychology, 89(1), 127–136. 10.1037/0021-9010.89.1.127

4. Anttila, H., Sullanmaa, J., & Pyhältö, K. (2021). Does it feel the same? Danish and Finnish social science and humanities doctoral students’ academic emotions. Frontiers in Education, 6, 758179.

5. Azen, R., & Budescu, D. V. (2003). The dominance analysis approach for comparing predictors in multiple regression. Psychological Methods, 8(2), 129.

6. Bagozzi, R. P. (1981). Evaluating structural equation models with unobservable variables and measurement error: A comment. Journal of Marketing Research, 18(3), 375–381.

7. Campbell, D. T., & Fiske, D. W. (1959). Convergent and discriminant validation by the multitrait-multimethod matrix. Psychological Bulletin, 56(2), 81.

8. Clark, R. A., Harden, S. L., & Johnson, W. B. (2000). Mentor relationships in clinical psychology doctoral training: Results of a national survey. Teaching of Psychology, 27(4), 262–268.

9. Danner, D., Aichholzer, J., & Rammstedt, B. (2015). Acquiescence in personality questionnaires: Relevance, domain specificity, and stability. Journal of Research in Personality, 57, 119–130.

10. Donnellan, M. B., Oswald, F. L., Baird, B. M., & Lucas, R. E. (2006). The mini-IPIP scales: Tiny-yet-effective measures of the Big Five factors of personality. Psychological Assessment, 18(2), 192.

11. Eby, L., & Allen, T. (2002). Further Investigation of Protégés’ Negative Mentoring Experiences: Patterns and Outcomes. Group & Organization Management, 27(4), 456–479.

12. Eby, L., Buits, M., Lockwood, A., & Simon, S. A. (2004). Protégés negative mentoring experiences: Construct development and nomological validation. Personnel Psychology, 57(2), 411–447.

13. Eby, L., McManus, S. E., Simon, S. A., & Russell, J. E. A. (2000). The Protege’s Perspective Regarding Negative Mentoring Experiences: The Development of a Taxonomy. Journal of Vocational Behavior, 57(1), 1–21.

14. Eby, L. T., Allen, T. D., Evans, S. C., Ng, T., & DuBois, D. L. (2008). Does mentoring matter? A multidisciplinary meta-analysis comparing mentored and non-mentored individuals. Journal of Vocational Behavior, 72(2), 254–267.

15. Eby, L. T., Butts, M. M., Durley, J., & Ragins, B. R. (2010). Are bad experiences stronger than good ones in mentoring relationships? Evidence from the protégé and mentor perspective. Journal of Vocational Behavior, 77(1), 81–92.

16. Eby, L. T. de T., Allen, T. D., Hoffman, B. J., Baranik, L. E., Sauer, J. B., Baldwin, S., Morrison, M. A., Kinkade, K. M., Maher, C. P., & Curtis, S. (2013). An interdisciplinary meta-analysis of the potential antecedents, correlates, and consequences of protégé perceptions of mentoring. Psychological Bulletin, 139(2), 441.

17. Eby, L. T., Durley, J. R., Evans, S. C., & Ragins, B. R. (2008). Mentors’ perceptions of negative mentoring experiences: Scale development and nomological validation. Journal of Applied Psychology, 93(2), 358.

18. Eby, L. T., & McManus, S. E. (2004). The protégé’s role in negative mentoring experiences. Journal of Vocational Behavior, 65(2), 255–275.

19. Eisenberger, R., Stinglhamber, F., Vandenberghe, C., Sucharski, I. L., & Rhoades, L. (2002). Perceived supervisor support: Contributions to perceived organizational support and employee retention. Journal of Applied Psychology, 87(3), 565.

20. Estrada, M., Zhi, Q., Nwankwo, E., & Gershon, R. (2019). The Influence of Social Supports on Graduate Student Persistence in Biomedical Fields. CBE—Life Sciences Education, 18(3), ar39.

21. Fagenson-Eland, E. A., Gayle, B. S., & Lankau, M. J. (2005). Seeing eye to eye: A dyadic investigation of the effect of relational demography on perceptions of mentoring activities. Career Development International, 10(6/7), 460–477.

22. Feeney, B. C., & Collins, N. L. (2015). A new look at social support: A theoretical perspective on thriving through relationships. Personality and Social Psychology Review, 19(2), 113–147.

23. Fleming, M., House, S., Hanson, V. S., Yu, L., Garbutt, J., McGee, R., Kroenke, K., Abedin, Z., & Rubio, D. M. (2013). The Mentoring Competency Assessment: Validation of a New Instrument to Evaluate Skills of Research Mentors. Academic Medicine, 88(7), 1002–1008.

24. Flesch, R. (1948). A new readability yardstick. Journal of Applied Psychology, 32(3), 221.

25. Floyd, F. J., & Widaman, K. F. (1995). Factor analysis in the development and refinement of clinical assessment instruments. Psychological Assessment, 7(3), 286.

26. Griffin, K. A., Stone, J., Dissassa, D. T., Hall, T. N., & Clarke, A. (2023). Surviving or flourishing: How relationships with principal investigators influence science graduate students’ wellness. Studies in Graduate and Postdoctoral Education, 14(1), 47–62.

27. Golde, C. M., & Dore, T. M. (2001). At Cross Purposes: What the Experiences of Today’s Doctoral Students Reveal about Doctoral Education.

28. Golino, H. F., & Epskamp, S. (2017). Exploratory graph analysis: A new approach for estimating the number of dimensions in psychological research. PloS One, 12(6), e0174035.

29. Golino, H., Shi, D., Christensen, A. P., Garrido, L. E., Nieto, M. D., Sadana, R., … & Martinez-Molina, A. (2020). Investigating the performance of exploratory graph analysis and traditional techniques to identify the number of latent factors: A simulation and tutorial. Psychological Methods, 25(3), 292.

30. Golino, H., & Christensen, A. (2024). EGAnet: Exploratory Graph Analysis–A Framework for Estimating the Number of Dimensions in Multivariate Data Using Network Psychometrics. (2025).

31. Handelsman, J., Pfund, C., Miller Lauffer, S., & Maidl Pribbenow, C. (2011). Entering mentoring: A seminar to train a new generation of scientists.

32. Harris, K. J., Kacmar, K. M., & Zivnuska, S. (2007). An investigation of abusive supervision as a predictor of performance and the meaning of work as a moderator of the relationship. *The Leadership Quarterly*, Destructive Leadership, 18(3), 252–263.

33. Harris, K., & Vazire, S. (2016). On friendship development and the Big Five personality traits. Social and Personality Psychology Compass, 10(11), 647–667.

34. Hibbing, M. V., Cawvey, M., Deol, R., Bloeser, A. J., & Mondak, J. J. (2019). The relationship between personality and response patterns on public opinion surveys: The big five, extreme response style, and acquiescence response style. International Journal of Public Opinion Research, 31(1), 161–177.

35. Hyun, S. H., Rogers, J. G., House, S. C., Sorkness, C. A., & Pfund, C. (2022). Revalidation of the Mentoring Competency Assessment to evaluate skills of research mentors: The MCA-21. Journal of Clinical and Translational Science, 6(1), e46.

36. Jiménez, M., Abad, F. J., Garcia-Garzon, E., Golino, H., Christensen, A. P., & Garrido, L. E. (2025). Dimensionality assessment in bifactor structures with multiple general factors: A network psychometrics approach. Psychological Methods, 30(4), 770.

37. John, O. P., & Srivastava, S. (1999). The Big-Five trait taxonomy: History, measurement, and theoretical perspectives.

38. Johnson, W. B. (2003). A framework for conceptualizing competence to mentor. Ethics & Behavior, 13(2), 127–151.

39. Johnson, W. B., & Huwe, J. M. (2002). Toward a typology of mentorship dysfunction in graduate school. Psychotherapy: Theory, Research, Practice, Training, 39(1), 44.

40. Kalbfleisch, P. J. (1997). Appeasing the mentor. Aggressive Behavior, 23(5), 389–403.

41. Kardash, C. M. (2000). Evaluation of undergraduate research experience: Perceptions of undergraduate interns and their faculty mentors. Journal of Educational Psychology, 92(1), 191.

42. Kincaid, J. P., Fishburne Jr, R. P., Rogers, R. L., & Chissom, B. S. (1975). Derivation of new readability formulas (automated readability index, fog count and flesch reading ease formula) for navy enlisted personnel.

43. Kram, K. E. (1985). Improving the mentoring process. Training & Development Journal.

44. Kram, K. E. (1988). Mentoring at work: Developmental relationships in organizational life. University Press of America.

45. Limeri, L. B., Asif, M. Z., Bridges, B. H. T., Esparza, D., Tuma, T. T., Sanders, D., Morrison, A. J., Rao, P., Harsh, J. A., Maltese, A. V., & Dolan, E. L. (2019). “Where’s My Mentor?!” Characterizing Negative Mentoring Experiences in Undergraduate Life Science Research. CBE—Life Sciences Education, 18(4), ar61.

46. Limeri, L. B., Carter, N. T., Hess, R. A., Tuma, T. T., Koscik, I., Morrison, A. J., Outlaw, B., Royston, K. S., Bridges, B. H. T., & Dolan, E. L. (2024). Development of the Mentoring in Undergraduate Research Survey. CBE—Life Sciences Education, 23(2), ar26.

47. Maher, M. A., Wofford, A. M., Roksa, J., & Feldon, D. F. (2020). Exploring early exits: Doctoral attrition in the biomedical sciences. *Journal of College Student Retention: Research*, Theory & Practice, 22(2), 205–226.

48. Mount, M. K., Barrick, M. R., & Stewart, G. L. (1998). Five-Factor Model of personality and Performance in Jobs Involving Interpersonal Interactions. Human Performance, 11(2–3), 145–165.

49. Nahm, A. Y., Rao, S. S., Solis-Galvan, L. E., & Ragu-Nathan, T. S. (2002). The Q-Sort Method: Assessing Reliability And Construct Validity Of Questionnaire Items At A Pre-Testing Stage. Journal of Modern Applied Statistical Methods, 1(1), 114–125.

50. National Academies of Sciences, Engineering, & Medicine. (2019). The Science of Effective Mentorship in STEMM.

51. Netemeyer, R. G., Boles, J. S., & McMurrian, R. (1996). Development and validation of work– family conflict and family–work conflict scales. Journal of Applied Psychology, 81(4), 400.

52. Nimon, K., Oswald, F., & Roberts, J. K. (2025). yhat: Interpreting Regression Effects (Version 2.0-5) [Computer software]. https://cran.r-project.org/web/packages/yhat/index.html

53. Noe, R. A. (1988). An investigation of the determinants of successful assigned mentoring relationships. Personnel Psychology, 41(3), 457–479.

54. Noy, S., & Ray, R. (2012). Graduate Students’ Perceptions of Their Advisors: Is There Systematic Disadvantage in Mentorship? The Journal of Higher Education, 83(6), 876– 914.

55. Porath, C. L., & Erez, A. (2009). Overlooked but not untouched: How rudeness reduces onlookers’ performance on routine and creative tasks. Organizational Behavior and Human Decision Processes, 109(1), 29–44.

56. Porath, C. L., & Pearson, C. M. (2010). The Cost of Bad Behavior. Organizational Dynamics, 39(1), 64–71.

57. R Core Team. (2021). R: A language and environment for statistical computing. R Foundation for Statistical Computing. *(No Title)*.

58. Ragins, B. R., Cotton, J. L., & Miller, J. S. (2000). Marginal Mentoring: The Effects of Type of Mentor, Quality of Relationship, and Program Design on Work and Career Attitudes. The Academy of Management Journal, 43(6), 1177–1194.

59. Reis, H. T., & Gable, S. L. (2015). Responsiveness. Current Opinion in Psychology, 1, 67–71.

60. Revelle, W. (2011). An overview of the psych package.

61. Rinker, T. (2015). Trinker/readability.

62. Rönkkö, M., & Cho, E. (2022). An Updated Guideline for Assessing Discriminant Validity. Organizational Research Methods, 25(1), 6–14.

63. Rosseel, Y. (2012). lavaan: An R Package for Structural Equation Modeling. Journal of Statistical Software, 48, 1–36.

64. Ruud, C. M., Saclarides, E. S., George-Jackson, C. E., & Lubienski, S. T. (2018). Tipping points: Doctoral students and consideration of departure. *Journal of College Student Retention: Research*, Theory & Practice, 20(3), 286–307.

65. Salmela-Aro, K., Kiuru, N., Leskinen, E., & Nurmi, J.-E. (2009). School Burnout Inventory (SBI): Reliability and Validity. European Journal of Psychological Assessment, 25(1), 48–57.

66. Scandura, T. A. (1998). Dysfunctional mentoring relationships and outcomes. Journal of Management, 24(3), 449–467.

67. Schlosser, L. Z., & Gelso, C. J. (2001). Measuring the working alliance in advisor–advisee relationships in graduate school. Journal of Counseling Psychology, 48(2), 157–167. (2001-00732-006).

68. Spitzer, R. L., Kroenke, K., Williams, J. B. W., & Löwe, B. (2006). A Brief Measure for Assessing Generalized Anxiety Disorder: The GAD-7. Archives of Internal Medicine, 166(10), 1092–1097.

69. Sverdlik, A., C. Hall, N., McAlpine, L., & Hubbard, K. (2018). The PhD Experience: A Review of the Factors Influencing Doctoral Students’ Completion, Achievement, and Well-Being. International Journal of Doctoral Studies, 13, 361–388.

70. Tenenbaum, H. R., Crosby, F. J., & Gliner, M. D. (2001). Mentoring relationships in graduate school. Journal of Vocational Behavior, 59(3), 326–341.

71. Tuma, T. T., Adams, J. D., Hultquist, B. C., & Dolan, E. L. (2021). The Dark Side of Development: A Systems Characterization of the Negative Mentoring Experiences of Doctoral Students. CBE—Life Sciences Education, 20(2), ar16.

72. Tuma, T. T., Fedesco, H. N., Rosenzweig, E. Q., Chen, X.-Y., & Dolan, E. L. (2025). Seeing isn’t believing? Mixed effects of a perspective-getting intervention to improve mentoring relationships for science doctoral students. CBE—Life Sciences Education, 24(4), ar44.

73. Ward, M. K., & Meade, A. W. (2023). Dealing with Careless Responding in Survey Data: Prevention, Identification, and Recommended Best Practices. Annual Review of Psychology, 74(1), 577–596.

74. Watson, D., Clark, L. A., & Tellegen, A. (1988). Development and validation of brief measures of positive and negative affect: The PANAS scales. Journal of Personality and Social Psychology, 54(6), 1063.

75. Welsh, E. T., & Diehn, E. W. (2018). Mentoring and gender: Perception is not reality. Career Development International, 23(4), 346–359.

76. Willis, G. B. (1999). Cognitive interviewing: A “how to” guide. Research Triangle Park, NC: Research Triangle Institute.

77. Zellars, K. L., Tepper, B. J., & Duffy, M. K. (2002). Abusive supervision and subordinates’ organizational citizenship behavior. Journal of Applied Psychology, 87, 1068–1076.

78. Zhao, C., Golde, C. M., & McCormick, A. C. (2007). More than a signature: How advisor choice and advisor behaviour affect doctoral student satisfaction. Journal of Further and Higher Education, 31(3), 263–281.

79. Zhong, S., & Maccalla, N. (2024). Measuring Faculty Mentoring Competency: Establishing the Validity of a Short Form. The Chronicle of Mentoring & Coaching, 8(2), 53–69.

