## Supplemental Materials for "Advancing the Conceptualization and Measurement of Mentorship: The Mentoring Experiences in Research & Graduate Education (MERGE) Scale"

**Table of Contents**


| Contents | Pages |
|---|---|
| Phase 2 Items | S2-S3 |
| Phase 2 Exploratory Graph Analyses and Exploratory Factor Analyses | S4-S5 |
| Phase 2 MERGE Item History | S6-S11 |
| Phase 2 Measurement Models | S12 |
| Phase 2 Dominance Analyses | S13 |
| Phase 3 Criterion-Related Validity Variables | S14-S15 |
| Phase 3 Items | S16-S18 |
| Phase 3 Measurement Models | S19-S20 |
| Phase 3 Incremental Validity Analyses | S21 |
| Phase 3 MCA-21 Facet-Level Dominance Analyses | S22 |
| References | S23-24 |

#### Phase 2 Items

**Graduate program support**
Eisenberger, R., Cummings, J., Armeli, S., & Lynch, P. (1997). Perceived organizational support, discretionary treatment, and job satisfaction. *Journal of Applied Psychology*, *82*(5), 812.

**Instructions:** Now please thinking about graduate program in general, not your mentor. Please indicate the extent to which you agree or disagree with each of these statements.
**Response Scale:** 1 = strongly disagree; 2 = disagree; 3 = slightly disagree; 4 = undecided; 5 = slightly agree; 6 = agree; 7 = strongly agree; 8 = prefer not to respond

1. My graduate program cares about my opinions.
2. My graduate program really cares about my well-being.
3. My graduate program strongly considers my goals and values.
4. Help is available from my graduate program when I have a problem.
5. My graduate program would forgive an honest mistake on my part.
6. If given the opportunity, my graduate program would take advantage of me. (R)
7. My graduate program shows very little concern for me. (R)
8. My graduate program is willing to help me if I need a special favor.

**Mentoring relationship quality & effectiveness**
Allen, T. D., & Eby, L. T. (2003). Relationship effectiveness for mentors: Factors associated with learning and quality. *Journal of Management*, *29*(4), 469-486.

**Instructions:** Please indicate the extent to which you agree or disagree with each of these statements.
**Response Scale:** 1 = strongly disagree; 2 = disagree; 3 = undecided; 4 = agree; 5 = strongly agree, 6 = Prefer not to respond

1. The mentoring relationship between my mentor and me is very effective.
2. I am very satisfied with the mentoring relationship my mentor and I have developed.
3. I am effectively utilized as mentee by my mentor.
4. My mentor and I enjoy a high-quality relationship.
5. Both my mentor and I benefit from our mentoring relationship.

**Personality**
Donnellan, M. B., Oswald, F. L., Baird, B. M., & Lucas, R. E. (2006). The mini-IPIP scales: tiny-yet-effective measures of the Big Five factors of personality. *Psychological Assessment*, *18*(2), 192.

**Instructions:** Please indicate how well each of the following statements describes you and your personality in general. Describe yourself as you typically are, not as you wish to be in the future.
**Response Scale:** 1 = very inaccurate; 2 = moderately inaccurate; 3 = neither accurate nor inaccurate; 4 = moderately accurate; 5 = very accurate; 6 = Prefer not to respond

**Extraversion**
1. I am the life of the party.
2. I don't talk a lot. (R)
3. I talk to a lot of different people at parties.
4. I keep in the background. (R)

**Agreeableness**
5. I sympathize with others' feelings.

6. I am not interested in other people’s problems. (R)
7. I feel others’ emotions.
8. I am not really interested in others. (R)

**Conscientiousness**

9. I get chores done right away.
10. I often forget to put things back in their proper place. (R)
11. I like order.
12. I make a mess of things. (R)

**Neuroticism**

13. I have frequent mood swings.
14. I am relaxed most of the time. (R)
15. I get upset easily.
16. I seldom feel blue. (R)

**Openness**

17. I have a vivid imagination.
18. I am not interested in abstract ideas. (R)
19. I have difficulty understanding abstract ideas.
20. I do not have a good imagination. (R)

**Positive & negative affect toward research**

Watson, D., Clark, L. A., & Tellegen, A. (1988). Development and validation of brief measures of positive and negative affect: the PANAS scales. *Journal of Personality and Social Psychology*, *54*(6), 1063.

**Instructions:** Now please thinking about graduate research in general, not your mentor. To what extent does your graduate research generally make you feel:
**Response Scale:** 1 = not at all; 2 = a little; 3 = a moderate amount; 4 = a lot; 5 = a great deal, 6 = Prefer not to respond

**Positive affect**

1. Alert
2. Inspired
3. Determined
4. Attentive
5. Active

**Negative affect**

6. Upset
7. Hostile
8. Ashamed
9. Nervous
10. Afraid

#### Phase 2 Exploratory Graph Analyses and Exploratory Factor Analyses

EGA is a modern network psychometrics approach for evaluating the internal structure that represents constructs as networks of items and uses algorithms to identify clusters of item covariation (i.e., dimensions - referred to as *communities*) within the network. One principal advantage of EGA over EFA is that EGA solutions do not rely on factor rotation; as such, decisions regarding internal structure are less subjective and more accurate compared to relying on solely traditional EFA procedures (Golino & Epskamp, 2017). Contemporary approaches to scale development emphasize the value of using both EGA and EFA because, while they differ in their underlying assumptions, they offer complementary insights for scale development and validation (Christensen & Golino, 2021).

We first used the unique variable analysis (UVA) function from the EGAnet package to identify empirically redundant items (Golino & Christensen, 2024). Redundant items, such as those with very similar wording, can produce spurious factors and lead to inadmissible factor solutions (Golino & Epskamp, 2017). As such, redundancies are important to remove prior to EGA and EFA. We reviewed each item pair that had a weighted topological overlap (wTO) >0.25 (Christensen et al., 2023), indicating empirical redundancy, and observed 12 redundant items. For each redundant pair, we retained the item with the greatest conceptual relevance and clearest wording.

We then used hierarchical EGA (Jiménez et al., 2025) to examine how the remaining items clustered together (i.e., the dimensional structure). EGA represents items as nodes in a network with connections that reflect item associations after accounting for other items in the network (Epskamp & Fried, 2018). Hierarchical EGA (hierEGA) is a new approach that allows for the detection of both specific item clusters (i.e., lower-order factors) and broader dimensions that encompass those clusters (i.e., higher order-factors). This approach is well-suited for our goals of understanding both dimensionality of the MERGE items and testing whether positive and negative mentoring experiences are empirically distinct constructs. We estimated a bootstrapped hierarchical EGA using the Louvain community detection algorithm with 500 parametric iterations to evaluate the stability of the number dimensions and network structure (Christensen et al., 2023). The initial hierarchical EGA returned a solution with 12 lower-order communities (95% C.I. [9.17, 14.83]) and 3 higher-order communities (95% CI [-.69, 2.69]); the wide confidence intervals indicated substantial variability in the structure across replications. Indeed, the typical lower-order structure (12 communities) emerged in only 25% of replications, and the typical higher-order structure (3 communities) emerged in only 23% of replications; in the majority of replications (63.6%), a single higher-order community emerged. Structural consistency - a measure of the extent to which a community was represented by the same set of items across replications - ranged from .17-1.00. In addition, average item stability - the extent to which the final set of representing a cluster were consistently assigned to that cluster across replications - ranged from .47-1.00. Item stability ranged from .21 to 1.00. Thus, although some clusters and items exhibited good psychometric properties, additional model refinement was needed to ensure the internal structure was stable and replicable.

To improve model stability, parsimony, and factor interpretability, we used the hierEGA results to inform a series of EFAs estimated using maximum likelihood robust (MLR) estimation with target rotation (i.e., the target matrix was specified according to the hierEGA-derived structure). Target rotation allowed us to evaluate whether the observed network-based dimensionality suggested by the hierEGA could be recovered as a latent factor structure, while still allowing the items to deviate from the target structure if empirically justified. We began by specifying an 11-factor EFA model based, which was targeted to match the hierarchical EGA lower-order structure with one exception. Specifically, we combined the two-item *Mentoring Satisfaction* community with the *Relationship Quality* community to avoid an under identified factor and because the *Mentoring Satisfaction* community demonstrated very low average item stability in the hierEGA (stability = .47) and its two items frequently clustered into *Relationship Quality*.

The initial EFA did not recover all 11 factors because some factors had no items loading onto them, while other comprised items from multiple communities. These findings indicate that the 11-factor solution was over-extracting, and that a more parsimonious solution was needed. We therefore sequentially combined factors and removed items that performed poorly across the EGA and EFA models. Specifically, we removed items when they (1) loaded weakly (<.30) on their target factor, (2) showed substantial or inconsistent cross-loadings, or (3) were not consistently assigned to the same dimension across the EGA and EFA. We also reassessed redundancy after each iteration as to ensure that retained items had UVA values < .25. We repeated this process until the remaining items demonstrated clear and consistent factor membership across the hierEGA and EFA models and the resulting dimensions were stable (i.e., items were consistently assigned to the same factor across replications) and interpretable (i.e., comprised coherent, conceptually distinct items). The hierEGA typical structure was six lower-order communities (95% CI 5.28, 6.72], with six lower-order communities emerging in 85.8% of replicates, and a single higher-order community (95% CI 1, 1], with a single higher-order community emerging in 100% of replicates. Across bootstrap replications, the six-factor hierEGA model demonstrated good structural stability, with average structural consistency estimates ranging from .88-1.00 and average item stability estimates ranging from .97 to 1.00. These results supported a stable six-factor structure with a single higher-order community.

#### Phase 2 MERGE Item History

| Item | Item Wording | Outcome | Rationale for Exclusion |
|---|---|---|---|
| MERGE1 | My mentor is willing to give me feedback on my research | Retained (Responsiveness) | |
| MERGE2 | My mentor advocates on my behalf | Excluded (UVA redundancy) | Redundant with MERGE3, wTO = .30 |
| MERGE3 | My mentor protects me from others who might cause me professional harm | Retained (Career Support) | |
| MERGE4 | My mentor has little interest in my career advancement | Retained (Career Support) | |
| MERGE5 | My mentor is reluctant to let me present my research at conferences | Excluded (Modification index) | Large modification index due to correlated uniqueness with MERGE4 (MI = 27.1); MERGE4 retained for the stronger primary loading |
| MERGE6 | My mentor offers useful advice for achieving my career goals | Retained (Career Support) | |
| MERGE7 | My mentor prioritizes publishing my research | Retained (Career Support) | |
| MERGE8 | My mentor helps me identify ways to network | Retained (Career Support) | |
| MERGE9 | My mentor makes sure I have sufficient funding to do my research | Excluded (Weak loading) | Maximum standardized loading .27 (Career Support, p=.002), below the .30 retention threshold |
| MERGE10 | My mentor helps me prepare for important milestones in my degree | Retained (Career Support) | |
| MERGE11 | My mentor says one thing and does another | Retained (Negative Mentoring Behaviors) | |
| MERGE12 | My mentor likes to stretch the truth in front of others | Excluded (Modification index) | Large modification index due to correlated uniqueness with MERGE11 (MI = 57.2); MERGE11 retained due to greater clarity |

| Item | Item Wording | Outcome | Rationale for Exclusion |
|---|---|---|---|
| MERGE13 | My mentor is only nice to me when they need me | Retained (Negative Mentoring Behaviors) | |
| MERGE14 | My mentor was nicer to me before I committed to work with them | Excluded (Weak loading) | Maximum standardized loading .26 (Negative, $p<.001$), below the .30 retention threshold |
| MERGE15 | My mentor lies to me | Excluded (Modification index) | Large modification indices with four items: MERGE13 (MI = 35.4), MERGE29 (MI = 28.6), MERGE11 (MI = 25.7), MERGE61 (MI = 22.3) |
| MERGE16 | My mentor intentionally misleads me | Excluded (UVA redundancy) | Redundant with MERGE15, wTO = .28; MERGE15 retained because it is less inferential |
| MERGE17 | My mentor is unfamiliar with my research topic | Excluded (UVA redundancy) | Redundant with MERGE18, wTO = .28; MERGE18 retained [though ultimately excluded] |
| MERGE18 | My mentor is unable to provide guidance on my research | Excluded (Modification index) | Large modification index due to correlated uniqueness with MERGE1 (MI = 17.8, EPC = 0.1); retained MERGE1 because more clearly worded |
| MERGE19 | My mentor lacks the technical knowledge to guide me on my research. | Excluded (UVA redundancy) | Redundant with MERGE18, wTO = .36; MERGE18 retained instead as more behaviorally worded |
| MERGE20 | My mentor has limited management expertise | Excluded (Weak loading) | Maximum standardized loading .24 (Career Support, $p=.002$), below the .30 retention threshold |
| MERGE21 | My mentor lacks mentoring competence | Excluded (UVA redundancy) | Redundant with MERGE20, wTO = .40 |
| MERGE22 | My mentor responds when I contact them | Retained (Responsiveness) | |
| MERGE23 | My mentor is too busy to meet with me | Excluded (Modification index) | Large modification index due to correlated uniqueness with MERGE26 (MI = 37.1) |
| MERGE24 | My mentor often forgets about my research progress | Retained (Responsiveness) | |
| MERGE25 | My mentor gives me their full attention when we meet | Retained (Responsiveness) | |

| Item | Item Wording | Outcome | Rationale for Exclusion |
|---|---|---|---|
| MERGE26 | My mentor's personal demands limit the time they have to mentor me | Retained (Responsiveness) | |
| MERGE27 | My mentor treats me the same as other mentees even though I have different needs | Excluded (EGA) | Unassigned in EGA; hierEGA would not run with it included due to lack of community assignment, so it was dropped before that step |
| MERGE28 | My mentor has an unpredictable personality | Retained (Negative Mentoring Behaviors) | |
| MERGE29 | My mentor is impatient | Retained (Negative Mentoring Behaviors) | |
| MERGE30 | My mentor tells me information about their personal life that makes me uncomfortable | Excluded (UVA redundancy) | Redundant with MERGE31, wTO = .27; MERGE30 judged more about the mentee's reaction than the mentor's behavior |
| MERGE31 | My mentor gives me advice on topics that are none of their business | Retained (Negative Mentoring Behaviors) | |
| MERGE32 | My mentor gets annoyed easily | Excluded (Heywood case) | Contributed to persistent inadmissible solutions (factor loadings > 1) |
| MERGE33 | My mentor thinks they are better than others | Retained (Negative Mentoring Behaviors) | |
| MERGE34 | My mentor is hot tempered | Retained (Negative Mentoring Behaviors) | |
| MERGE35 | My mentor does not take responsibility for their mistakes | Retained (Negative Mentoring Behaviors) | |
| MERGE36 | My mentor and I think different things are important in life | Excluded (Weak loading) | Loaded < .30 on its target factor |
| MERGE37 | My mentor and I have incompatible personalities | Retained (Relationship Strain) | |

| Item | Item Wording | Outcome | Rationale for Exclusion |
|---|---|---|---|
| MERGE38 | My mentor has no respect for my career goals | Excluded (Network instability) | Instability across Career Support and Negative Mentoring Behaviors due to blended content |
| MERGE39 | My mentor's personality works well with mine | Excluded (Cross-loading) | Instability across Relationship Strain and Relationship Quality due to blended content |
| MERGE40 | My mentor and I have similar work styles | Excluded (Cross-loading) | Instability across Relationship Strain and Relationship Quality due to blended content |
| MERGE41 | My mentor and I argue based on our differences in values. | Excluded (Network instability) | Instability across Relationship Quality and Negative Mentoring Behaviors due to blended content |
| MERGE42 | The mentoring relationship between my mentor and I is very effective. | Retained (Relationship Quality) | |
| MERGE43 | I am very satisfied with the mentoring relationship my mentor and I have developed. | Excluded (Modification index) | Large modification index due to correlated uniqueness with MERGE42 (MI = 57.6); MERGE42 retained |
| MERGE44 | I am effectively utilized as a mentee by my mentor. | Retained (Relationship Quality) | |
| MERGE45 | My mentor and I enjoy a high-quality relationship. | Retained (Relationship Quality) | |
| MERGE46 | Both my mentor and I benefit from our mentoring relationship. | Retained (Relationship Quality) | |
| MERGE47 | My mentor encourages me | Retained (Psychosocial Support) | |
| MERGE48 | My mentor values me as a person | Excluded (Modification index) | Large modification index due to correlated uniqueness with MERGE47 (MI = 34.8, EPC = 0.1); MERGE47 retained |
| MERGE49 | My mentor checks in about my well-being | Retained (Psychosocial Support) | |

| Item | Item Wording | Outcome | Rationale for Exclusion |
|---|---|---|---|
| MERGE50 | My mentor is a role model for me. | Excluded (Weak loading) | Maximum standardized loading .27 (Relationship Quality, p<.001), below the .30 retention threshold |
| MERGE51 | My mentor makes me feel accepted | Retained (Psychosocial Support) | |
| MERGE52 | My mentor empathizes when I am struggling | Retained (Psychosocial Support) | |
| MERGE53 | My mentor is understanding when I experience difficulties | Excluded (UVA redundancy) | Redundant with MERGE52 wTO = .29; MERGE52 retained instead as more clearly about behavior than internal state |
| MERGE54 | My mentor tells me when they think I have done a good job | Retained (Psychosocial Support) | |
| MERGE55 | My mentor and I do not like each other | Retained (Relationship Strain) | |
| MERGE56 | My mentor and I have a difficult relationship | Retained (Relationship Strain) | |
| MERGE57 | My mentor and I can talk about things other than work tasks. | Excluded (Cross-loading) | Instability across Psychosocial Support and Relationship Strain due to shared content |
| MERGE58 | My mentor and I have a tense relationship | Retained (Relationship Strain) | |
| MERGE59 | My mentor encourages competition between lab members | Excluded (Weak loading) | Maximum standardized loading .27 (Negative Mentoring Behaviors, p<.001), below the .30 retention threshold |
| MERGE60 | My mentor makes me do tasks not related to our lab's research | Excluded (Weak loading) | Maximum standardized loading -.25 (Responsiveness, p<.001), below the .30 retention threshold |
| MERGE61 | My mentor threatens me | Retained (Negative Mentoring Behaviors) | |

| Item | Item Wording | Outcome | Rationale for Exclusion |
|---|---|---|---|
| MERGE62 | My mentor calls me insulting names | Excluded (Modification index) | Large modification index due to correlated uniqueness with MERGE61 (MI = 63.1, EPC = 0.1); MERGE61 retained |
| MERGE63 | My mentor yells at me | Excluded (Near-redundancy) | wTO with MERGE62= .25 |
| MERGE64 | My mentor takes their stress out on me | Retained (Negative Mentoring Behaviors) | |
| MERGE65 | My mentor wants me to become independent before I am ready. | Excluded (Weak loading) | Strongest loading across all 6 factors was .28 (Relationship Quality, p=.019); loaded only .26 on its own assigned Responsiveness target |
| MERGE66 | My mentor keeps up with my research progress | Retained (Responsiveness) | |
| MERGE67 | My mentor is not involved enough in my research | Retained (Responsiveness) | |
| MERGE68 | My mentor is too controlling | Excluded (UVA redundancy) | Redundant with MERGE69, wTO = .27; MERGE69 retained |
| MERGE69 | My mentor micromanages me | Retained (Negative Mentoring Behaviors) | |
| MERGE70 | My mentor tries to get too involved in the day-to-day aspects of my research | Excluded (UVA redundancy) | Redundant with MERGE69, wTO = .26; MERGE69 retained |
| MERGE71 | My mentor likes to approve of minor trivial decisions | Excluded (UVA redundancy) | Redundant with MERGE69, wTO = .38; MERGE69 retained |
| MERGE72 | My mentor monitors my work too closely | Excluded (UVA redundancy) | Redundant with both MERGE71 (wTO = .31) and MERGE69 (wTO = .26); MERGE69 retained |

*Of 72 candidate items entered into the item-reduction analysis, 37 were retained in the final 6-factor structure and 35 were excluded via redundancy screening (unique-variable analysis, UVA), network-structure checks (EGA/hierEGA/bootEGA), weak or cross-loadings in exploratory factor analysis, or modification-index-driven review of semantically overlapping items. The label in parentheses in the Outcome column indicates the type of evidence behind each exclusion; the Rationale column gives the specific statistic (wTO, standardized loading, modification index, or bootEGA community-placement percentage) that justified it. MI = modification index; EPC = expected parameter change. MERGE items 46-46 are adapted from Allen, T. D., & Eby, L. T. (2003). Relationship effectiveness for mentors: Factors associated with learning and quality. Journal of Management, 29(4), 469-486.*

#### Phase 2 Measurement Models

| Measure | Model | $\chi^2$ (*df*) | CFI | TLI | RMSEA | SRMR | *N* | Notes |
|---|---|---|---|---|---|---|---|---|
| Graduate program support | Original | 144.15 (20) | 0.951 | 0.932 | 0.122 | 0.040 | 530 | Acceptable fit as published; largest modification index is pos6~~pos7 (both reverse-worded items) |
| Graduate Program Support | Single-Factor CFA with Correlated Uniqueness | 108.92 (19) | 0.965 | 0.948 | 0.106 | 0.037 | 530 | pos6~~pos7 freed |
| Positive & negative affect towards graduate research | Original | 321.30 (34) | 0.867 | 0.823 | 0.132 | 0.100 | 530 | Poor fit; posaff1 ("Alert") loads only .236 and cross-loads onto Negative Affect (MI = .469); negaff4/negaff5 share anxiety-specific residual content |
| Positive & negative affect towards graduate research | Five-Factor CFA with 2 Dropped Items | 98.11 (19) | 0.955 | 0.934 | 0.093 | 0.043 | 530 | posaff1 and negaff4 dropped |
| Big Five Personality (mini-IPIP) | Five-Factor CFA | 615.50 (160) | 0.857 | 0.831 | 0.076 | 0.066 | 530 | Poor-to-modest fit; largest MIs are within-facet reverse-keyed-pair residual correlations (method-variance pattern) |
| Big Five Personality (mini-IPIP) | Five-Factor CFA with Correlated Uniquenesses | 445.01 (155) | 0.910 | 0.890 | 0.061 | 0.064 | 530 | 5 reverse-keyed residual covariances added (extrav2~~extrav4, agree2~~agree4, consci2~~consci4, neuro2~~neuro4, intel2~~intel3) |
| All Measures | EFA (MERGE) + CFA | 3502.49 (2303) | 0.955 | 0.949 | 0.032 | 0.040 | 530 | MERGE (6-factor target-rotation EFA) + Graduate Program Support + PANAS(2) + Big Five(5) |

*Chi-square is the MLR-scaled test statistic; CFI, TLI, and RMSEA are robust variants. Grey shading indicates the final measurement model that was used in the substantive analyses.*

#### Phase 2 Dominance Analyses

We conducted dominance analyses to examine the relative importance of each MERGE factor for explaining variance in each criterion variable, after accounting for the overlap in variance explained by the other factors. Dominance analysis is a variance decomposition technique used to estimate the relative importance of correlated predictors by evaluating their contribution to explained variance ($R^2$) across all possible subset regression models. This approach is particularly useful in the presence of multicollinearity among predictors, as it provides a more stable and interpretable index of predictor importance than traditional regression coefficients. Whereas bivariate approaches used indicate whether constructs are associated in theoretically expected directions, they do not provide information about the unique explanatory contribution of each construct when predictors share overlapping variance. Dominance analysis addresses this limitation by partitioning shared and unique variance across predictors and evaluating their relative importance across all model specifications.

We conducted the dominance analyses following the approach described by Azen and Budescu (2003). We extracted latent factor scores from the CFA measurement model using regression-based estimates in lavaan. Dominance analyses were based on linear regression models that regressed the criterion measures (i.e., graduate program support, negative affect toward research, positive affect toward research) on the set of MERGE factors. We extracted the general dominance weights as the primary index of predictor importance because they reflect each predictor's average contribution to the total explained variance ($R^2$) across all models. They also provide an estimate of how much each MERGE factor uniquely contributes to predicting each criterion variable when the shared variance among MERGE factors is taken into account. Thus, $R^2$ represents the total proportion of variance in the outcome explained by the MERGE, whereas dominance weights represent each factor's proportional contribution to that explained variance.

#### Phase 3 Criterion-Related Validity Variables

***Research self-efficacy***

Self-efficacy refers to an individual's context-dependent belief in their ability to successfully complete a task or complete a specific goal. A doctoral student's belief about their scientific research self-efficacy is an important predictor of their goal-setting, effort, persistence, performance, and wellbeing (Choi et al., 2012; Pajares, 1996; Sitzmann & Yeo, 2013). Bandura's (1997) Social Cognitive Theory maintains that self-efficacy beliefs are derived from four types of experiential sources: mastery experiences, vicarious experiences, social persuasion, and task specific emotional. Social persuasion is particularly relevant to doctoral student mentoring because mentors can provide feedback, encouragement, and evaluations that shape students' beliefs about their research capabilities. Doctoral students' relationships with faculty and their mentoring experiences have been linked to their self-efficacy beliefs (Bishop & Bieschke, 1998; Ferguson & McHenry-Sorber, 2025; Tuma et al., 2025).

Mentoring experiences may strengthen or undermine students' research self-efficacy. Mentors who couple increased research independence with support may help build student confidence in their ability to successfully perform research tasks. Conversely, mentors who provide inadequate guidance, provide overly harsh feedback, or are unavailable to help students navigate research challenges may limit students' accomplishment of mastery experiences and thus their research self-efficacy. Overly discouraging criticism or failure to recognize students' research contributions may function as negative social persuasion by communicating doubts about students' research abilities. We hypothesized that supportive mentoring experiences would be positively associated with research self-efficacy, and negative mentoring experiences would be negatively associated with research self-efficacy.

***Burnout***

Burnout is a state of physical, mental, or emotional exhaustion resulting from prolonged exposure to stress (Maslach, 2011). According to the Job Demands-Resources Model, burnout is more likely to develop when demands are high and the resources available to meet those demands are insufficient (Bakker & Demerouti, 2007). For doctoral students, mentoring relationships may be an important source of resources for managing the demands of graduate education. For example, supportive mentoring experiences may provide guidance, feedback, and encouragement that help doctoral students manage the demands of doctoral research and training. Negative mentoring experiences may reduce access to these resources while introducing additional interpersonal demands. Indeed, doctoral students who report sufficient support from their supervisors are less likely to report burnout than students who reported insufficient support (Pyhältö et al., 2017). Doctoral students who report receiving sufficient supervisory support are also at lower risk of burnout symptoms (Tikkanen et al., 2025). We hypothesized that supportive mentoring experiences would be negatively associated with student reports of burnout, and negative mentoring experiences would be positively associated.

***Work-family conflict***

Work-family conflict refers to a form of inter-role conflict in which the demands from work and family domains are incompatible, making it difficult to meet responsibilities in both areas (Greenhaus & Beutell, 1985). Work-family conflict can occur when work-related demands interfere with responsibilities outside of work (i.e., work interference with family) or when family demands interfere with work responsibilities (i.e., family interference with work). We focused on work interference with family, operationalized as the extent to which doctoral students' research-related responsibilities interfered with their personal lives. This focus reflects the substantial time, effort, and investment that doctoral research can require and the potential for these demands to extend into students' lives outside of research.

Mentoring experiences may be relevant to work-family conflict because mentors can provide psychological and instrumental resources that help students manage the demands of doctoral training

(Cohen & Wills, 1985; House et al., 1988). When individuals have fewer resources to manage demanding roles, this may interfere with home and family responsibilities (Demerouti et al., 2001). Evidence from the work-family literature suggests that mentoring and supervisor support are associated with lower work-family conflict (Kossek et al., 2011; Nielson et al., 2001). We hypothesized that supportive mentoring experiences would be negatively associated with student reports of work-family conflict, and negative mentoring experiences would be positively associated.

***Anxiety***

Anxiety refers to a state characterized by excessive fear or worry and related cognitive, emotional, and physiological symptoms that can interfere with daily functioning (APA, 2022). Doctoral training involves academic, research, and interpersonal demands that may contribute to students' anxiety. Support from mentors may help students manage the challenges they encounter while conducting research, while negative mentoring experiences may contribute to or exacerbate these challenges. Some estimates suggest that anxiety may be common among graduate students, with over 40% reporting moderate-to-severe anxiety symptoms (Evans et al., 2018). Results from this same sample suggested that positive faculty mentoring relationships were associated with lower symptoms of anxiety (Evans et al., 2018). Among doctoral students specifically, more positive mentoring relationships were also associated with lower anxiety symptoms (Liu et al., 2019). We hypothesized that supportive mentoring experiences would be negatively associated with student reports of anxiety, and negative mentoring experiences would be positively associated.

***Career intentions***

Mentors can provide guidance and support that may help students explore and evaluate potential career paths (Allen et al., 2004). Doctoral students who report greater mentoring support also report greater academic career self-efficacy and, indirectly, interest in pursuing an academic career (Curtin et al., 2016). Research on life science doctoral students has similarly linked mentoring support with career preparedness (Chang et al., 2021). We hypothesized that supportive mentoring experiences to be positively related to students' career intentions, and negative mentoring experiences would be negatively associated.

#### Phase 3 Items

**Research self-efficacy**

Kardash, C. M. (2000). Evaluation of undergraduate research experience: Perceptions of undergraduate interns and their faculty mentors. *Journal of Educational Psychology*, *92*(1), 191.

**Instructions:** Please indicate your confidence in your ability to perform the following tasks.
**Response Scale:** 1 = not at all confident; 2 = not very confident; 3 = somewhat confident; 4 = fairly confident; 5 = very confident 6 = Prefer not to respond

*Note.* One item from the original scale ("identify a specific research question for investigation based on the research in your field") was inadvertently omitted from the survey.

1. Understand contemporary concepts in your field.
2. Make use of the primary scientific research literature in your field (e.g., journal articles).
3. Formulate a research hypothesis based on a specific question.
4. Design an experiment or theoretical test of the hypothesis.
5. Understand the importance of "controls" in research.
6. Observe and collect data.
7. Statistically analyze data.
8. Interpret data by relating results to the original hypothesis.
9. Reformulate your original research hypothesis (as appropriate).
10. Relate the results to the "bigger picture" in your field.
11. Orally communicate the results of research projects.
12. Write a research paper for publication.
13. Think independently.

**Career intentions**

Estrada, M., Zhi, Q., Nwankwo, E., & Gershon, R. (2019). The influence of social supports on graduate student persistence in biomedical fields. *CBE—Life Sciences Education*, *18*(3), ar39.

**Instructions:** Please indicate the extent to which you agree with the following statements.
**Response Scale:** 1 = not at all likely; 2 = a little bit likely; 3 = somewhat likely; 4 = very likely; 5 = absolutely likely, 6 = Prefer not to respond

1. To what extent do you intend to pursue a science-related research career?
2. To what extent do you intend to pursue a career in the same field as your graduate research?
3. To what extent do you intend to pursue a career in which you publish scientific findings in peer-reviewed journals?
4. To what extent do you intend to pursue a career in which you write grant applications for funding?

**Work-family conflict**

Netemeyer, R. G., Boles, J. S., & McMurrian, R. (1996). Development and validation of work–family conflict and family–work conflict scales. *Journal of Applied Psychology*, 81(4), 400.

**Instructions:** Please indicate the extent to which you agree with the following statements.
**Response Scale:** 1 = strongly disagree; 2 = disagree; 3 = undecided; 4 = agree; 5 = strongly agree, 6 = Prefer not to respond

1. The demands of my research interferes with my home/family life.
2. The amount of time my research takes up makes it difficult to fulfill home/family responsibilities.
3. Things I want to do at home do not get done because of the demands research puts on me.
4. My research produces strain that makes it difficult to fulfill my home/family duties.
5. Due to research-related duties, I have to make changes to my plans for home/family activities.

**<u>Burnout</u>**
Salmela-Aro, K., Kiuru, N., Leskinen, E., & Nurmi, J. E. (2009). School burnout inventory (SBI) reliability and validity. *European Journal of Psychological Assessment*, *25*(1), 48-57.

**Instructions:** Please indicate the extent to which you agree with the following statements.
**Response Scale:** 1 = completely disagree; 2 = partly disagree; 3 = disagree; 4 = partly agree; 5 = agree, 6 = completely agree, 7 = Prefer not to respond

1. I feel overwhelmed by my graduate program.
2. I feel a lack of motivation in my graduate program and often think of giving up.
3. I often have feelings of inadequacy in my graduate program.
4. I often sleep badly because of matters related to my graduate program.
5. I feel that I am losing interest in my graduate program.
6. I'm continually wondering whether my graduate program has any meaning.
7. I brood over matters related to my graduate program a lot during my free time.
8. I used to have higher expectations of my graduate program that I do now.
9. The pressure of my graduate program causes me problems in my close relationships with others.

**<u>Anxiety</u>**
Spitzer, R. L., Kroenke, K., Williams, J. B., & Löwe, B. (2006). A brief measure for assessing generalized anxiety disorder: the GAD-7. *Archives of Internal Medicine*, 166(10), 1092-1097.

**Instructions:** Over the last two weeks, how often have you been bothered by the following problems?
**Response Scale:** 1 = not at all; 2 = several days; 3 = more than half the days; 4 = nearly every day; 5 = Prefer not to respond

1. Feeling nervous, anxious, or on edge.
2. Not being able to stop or control worrying.
3. Worrying too much about different things.
4. Having trouble relaxing.
5. Being so restless that it is hard to sit still.
6. Becoming easily annoyed or irritable.
7. Feeling afraid as if something awful might happen.

**<u>Revised Mentoring Competency Assessment</u>**
Hyun, S. H., Rogers, J. G., House, S. C., Sorkness, C. A., & Pfund, C. (2022). Revalidation of the mentoring competency assessment to evaluate skills of research mentors: the MCA-21. *Journal of Clinical and Translational Science*, *6*(1), e46.

**Instructions:** Please rate how skilled your mentor is in the following areas.

**Response Scale:** 1 = not at all skilled; 2 = slightly skilled; 3 = somewhat skilled; 4 = neither skilled or unskilled; 5 = moderately skilled; 6 = very skilled; 7 = extremely skilled; 8 = Prefer not to respond

**Maintaining effective communication**

1. Active listening.
2. Providing constructive feedback.
3. Establishing a relationship based on trust.
4. Identifying and accommodating different communication styles.

**Aligning expectations**

5. Working with you to set clear expectations of the mentoring relationship.
6. Aligning their expectations with you.
7. Accurately estimating your ability to conduct research.
8. Employing strategies to enhance your knowledge and abilities.

**Assessing understanding**

9. Working with you to set research goals.
10. Helping you develop strategies to meet goals.
11. Accurately estimating your level of scientific knowledge.

**Fostering independence**

12. Stimulating your creativity.
13. Acknowledging your professional contributions.
14. Helping you negotiate a path to professional independence.

**Addressing diversity**

15. Considering how your personal and professional differences may impact expectations.
16. Taking into account the biases and prejudices they bring to the mentoring relationship.
17. Working effectively with mentees whose personal background is different from their own.

**Promoting professional development**

18. Helping you network effectively.
19. Helping you set career goals.
20. Helping you balance work with your personal life.
21. Helping you acquire resources.

**Phase 3 Measurement Models**

| Measure | Model | χ² (*df*) | CFI | TLI | RMSEA | SRMR | *N* | Notes |
|---|---|---|---|---|---|---|---|---|
| Work-Family Conflict | Single-Factor CFA | 26.00 (5) | 0.990 | 0.979 | 0.096 | 0.011 | 749 | |
| Anxiety | Single-Factor CFA | 79.71 (14) | 0.978 | 0.967 | 0.094 | 0.025 | 749 | |
| Career Intentions | Single-Factor CFA | 46.26 (2) | 0.955 | 0.866 | 0.198 | 0.047 | 749 | |
| Career Intentions | Single-Factor CFA with Uniqueness Covariance | 2.95 (1) | 0.998 | 0.990 | 0.054 | 0.008 | 749 | careerint3~~careerint4 freed |
| Burnout | Second-Order CFA with 3 First-Order Factors | Did not converge | -- | -- | -- | -- | 749 | Inadequacy factor is empirically underidentified (2 items). |
| Burnout | Single-Factor CFA with Item Parcels | 0.00 (0) | 1.000 | 1.000 | 0.000 | 0.000 | 749 | Single-factor model with 3 item-to-construct-balanced (mixed) parcels |
| Research Self-Efficacy | Single-Factor CFA | 390.93 (65) | 0.896 | 0.875 | 0.093 | 0.048 | 749 | |
| Research Self-Efficacy | Single-Factor CFA with Item Parcels | 0.00 (0) | 1.000 | 1.000 | 0.000 | 0.000 | 749 | Single-factor model with 3 item-to-construct-balanced (mixed) parcels |
| Mentor Competence Assessment (MCA-21) | Six-Factor CFA | 830.89 (174) | 0.944 | 0.932 | 0.087 | 0.034 | 749 | Strong interfactor intercorrelations (> .90) raise discriminant-validity concerns |

| Measure | Model | $\chi^2$ (*df*) | CFI | TLI | RMSEA | SRMR | *N* | Notes |
|---|---|---|---|---|---|---|---|---|
| Mentor Competence Assessment (MCA-21) | Second-Order CFA with 6 First-Order Factors | 1003.20 (183) | 0.930 | 0.920 | 0.095 | 0.037 | 749 | |
| All Measures | EFA (MERGE) + CFA | 6345.21 (2852) | 0.934 | 0.927 | 0.043 | 0.036 | 749 | MERGE (6-factor target-rotation EFA, 37-item sup11-retained structure) + Work-Family Conflict + Anxiety + Career Intentions + Burnout(parcels) + Self-Efficacy(parcels) + MCA(6-facet+2nd-order) |

*Chi-square is the MLR-scaled test statistic; CFI, TLI, and RMSEA are robust variants. Grey shading indicates the final measurement model that was used in the substantive analyses.*

**Phase 3 Incremental Validity Analyses**

We examined the incremental validity of the MERGE in explaining additional variance in theoretically relevant outcomes over and above the revised mentoring competence assessment (i.e., MCA-21) using hierarchical multiple regression (Hyun et al., 2022). We represented the MCA-21 as a single second-order total score rather than modeling its individual facets because the MCA-21 factors were extremely highly correlated (latent correlations = .79 to .95), making it difficult to distinguish their unique contributions.

For each outcome, we first estimated a model containing the MCA-21 score and then added the six MERGE factors simultaneously. We also tested the reverse comparison by adding the MCA-21 to a model that already included the six MERGE factors. We then examined the change in adjusted $R^2$ to determine the additional variance explained for each outcome. We also used nested-model $F$ tests to determine whether the increase in explained variance was statistically significant, and 95% bootstrap confidence intervals based on 1,000 replications to quantify uncertainty around the estimating increase and determine whether the interval included zero.

The MERGE demonstrated incremental validity beyond the MCA-21 for four of the five outcomes examined (Table 7). The MERGE accounted for unique variance in the prediction of work-family conflict over and above the MCA-21 (Δadj. $R^2 = .03$, $p < .001$, 95% CI [.01, .07]). In contrast, the MCA-21 did not account for unique variance in the prediction of work-family conflict over the MERGE (Δadj. $R^2 = .00$, $p = .124$, 95% CI [-.00, .01]). The MERGE also accounted for additional variance in the prediction of anxiety over the MCA-21 (Δadj. $R^2 = .03$, $p < .001$, 95% CI [0.01, .07]).), while the MCA-21 did not account for unique variance in anxiety over the MERGE (Δadj. $R^2 = -.00$, $p = .566$, 95% CI [-.00, .01]). Neither the MERGE nor the MCA-21 explained meaningful unique variance in career intentions over the other (MERGE over MCA-21: Δadj. $R^2 = -.00$, $p = .425$, 95% CI [-.00, .03]; MCA-21 over MERGE: Δadj. $R^2 = .00$, $p = .026$, 95% CI [-.00, .02]). The MERGE also accounted for unique variance in the prediction of burnout over the MCA-21 (Δadj. $R^2 = .03$, $p < .001$, 95% CI [.02, .07]), but the MCA-21 did not explain burnout beyond the MERGE (Δadj. $R^2 = .00$, $p = .013$, 95% CI [-.00, .02]). Finally, the MERGE accounted for unique variance in self-efficacy over the MCA-21 (Δadj. $R^2 = .04$, $p < .001$, 95% CI [.02, .08]). However, the MCA-21 also accounted for unique variance in self-efficacy over the MERGE (Δadj. $R^2 = .02$, $p < .001$, 95% CI [.00, .04]), but less than the MERGE.

### Phase 3 MCA-21 Facet-Level Dominance Analyses

| Predictor | Outcome | | | | | | | | | | | | | | |
|---|---|---|---|---|---|---|---|---|---|---|---|---|---|---|---|
| | Work-Family Conflict | | | Anxiety | | | Career Intentions | | | Burnout | | | Self-Efficacy | | |
| | β | Dom | $sr^2$ | β | Dom | $sr^2$ | β | Dom | $sr^2$ | β | Dom | $sr^2$ | β | Dom | $sr^2$ |
| Responsiveness | -0.02 | 0.01 | 0.00 | -0.02 | 0.01 | 0.00 | 0.03 | 0.01 | 0.00 | -0.07 | 0.02 | 0.00 | -0.08 | 0.00 | 0.00 |
| Career Support | 0.06 | 0.01 | 0.00 | 0.24* | 0.01 | 0.01 | 0.17 | **0.01** | 0.00 | 0.11 | 0.02 | 0.00 | 0.13 | 0.01 | 0.00 |
| Conflict | 0.04 | 0.01 | 0.00 | 0.03 | 0.01 | 0.00 | -0.02 | 0.01 | 0.00 | 0.05 | 0.03 | 0.00 | 0.09 | 0.01 | 0.00 |
| Psychosocial Support | -0.30* | 0.02 | 0.01 | -0.04 | 0.01 | 0.00 | -0.05 | 0.01 | 0.00 | 0.03 | 0.03 | 0.00 | -0.32* | 0.01 | 0.01 |
| Relationship Quality | 0.18 | 0.01 | 0.00 | -0.22 | 0.02 | 0.00 | -0.01 | 0.01 | 0.00 | -0.38* | **0.04** | 0.01 | 0.42* | **0.02** | 0.02 |
| Negative | 0.21* | **0.02** | 0.01 | 0.30* | **0.03** | 0.02 | -0.03 | 0.01 | 0.00 | 0.13 | 0.03 | 0.00 | 0.15 | 0.01 | 0.00 |
| Effective Communication | 0.37* | 0.01 | 0.01 | 0.42* | 0.01 | 0.01 | -0.10 | 0.01 | 0.00 | 0.46* | 0.03 | 0.01 | -0.28 | 0.01 | 0.00 |
| Aligning Expectations | -0.59* | 0.01 | 0.01 | -0.08 | 0.01 | 0.00 | 0.42 | 0.01 | 0.00 | -0.33 | 0.03 | 0.00 | 0.18 | 0.01 | 0.00 |
| Assessing Understanding | 0.35 | 0.01 | 0.00 | -0.00 | 0.01 | 0.00 | -0.20 | 0.01 | 0.00 | 0.09 | 0.02 | 0.00 | 0.20 | 0.02 | 0.00 |
| Fostering Independence | 0.24 | 0.01 | 0.00 | -0.11 | 0.01 | 0.00 | 0.20 | 0.01 | 0.00 | -0.04 | 0.03 | 0.00 | 0.50* | 0.02 | 0.01 |
| Addressing Diversity | -0.14 | 0.01 | 0.00 | -0.10 | 0.01 | 0.00 | -0.19 | 0.01 | 0.00 | -0.15 | 0.03 | 0.00 | 0.10 | 0.01 | 0.00 |
| Promoting Professional Development | -0.29 | 0.01 | 0.00 | -0.13 | 0.01 | 0.00 | 0.03 | 0.01 | 0.00 | -0.14 | 0.03 | 0.00 | -0.40* | 0.01 | 0.01 |
| ***$R^2$*** | | **0.16** | | | **0.14** | | | **0.13** | | | **0.34** | | | **0.12** | |

*Note. N = 747-749 (Phase 3, varies slightly by outcome). Predictors are the 6 MERGE factors (rows 1-6) and MCA-21's 6 facets. Beta (β) is the standardized OLS coefficient from a simultaneous regression of the outcome on all 12 predictors; * $p < .05$. Dom is the general dominance weight (each predictor's marginal contribution to $R^2$ across every possible subset of the other predictors, summing to the model $R^2$ reported in the bottom row for each outcome). $sr^2$ is the semi-partial $R^2$ (the drop in model $R^2$ if that predictor alone were removed, holding the other 11 in). Bolded Dominance values indicate the single most dominant predictor overall for that outcome (across both blocks).*

#### References


Allen, T. D., Eby, L. T., Poteet, M. L., Lentz, E., & Lima, L. (2004). Career Benefits Associated With Mentoring for Proteges: A Meta-Analysis. *Journal of Applied Psychology*, *89*(1), 127–136. https://doi.org/10.1037/0021-9010.89.1.127

American Psychological Association. (2022). *Anxiety*. https://www.apa.org/topics/anxiety

Azen, R., & Budescu, D. V. (2003). The dominance analysis approach for comparing predictors in multiple regression. *Psychological Methods*, *8*(2), 129.

Bakker, A. B., & Demerouti, E. (2007). The job demands-resources model: State of the art. *Journal of Managerial Psychology*, *22*(3), 309–328.

Bandura, A. (1997). *Self-Efficacy: The Exercise of Control, Nueva York, NH Freeman*.

Bishop, R. M., & Bieschke, K. J. (1998). Applying social cognitive theory to interest in research among counseling psychology doctoral students: A path analysis. *Journal of Counseling Psychology*, *45*(2), 182.

Chang, C.-N., Patterson, C. A., Vanderford, N. L., & Evans, T. M. (2021). Modeling individual development plans, mentoring support, and career preparedness relationships among Doctor of Philosophy (Ph. D.) trainees in the life sciences. *F1000Research*, *10*, 626.

Choi, B. Y., Park, H., Yang, E., Lee, S. K., Lee, Y., & Lee, S. M. (2012). Understanding Career Decision Self-Efficacy: A Meta-Analytic Approach. *Journal of Career Development*, *39*(5), 443–460. https://doi.org/10.1177/0894845311398042

Christensen, A. P., Garrido, L. E., & Golino, H. (2023). Unique Variable Analysis: A Network Psychometrics Method to Detect Local Dependence. *Multivariate Behavioral Research*, *58*(6), 1165–1182. https://doi.org/10.1080/00273171.2023.2194606

Christensen, A. P., & Golino, H. (2021). On the equivalency of factor and network loadings. *Behavior Research Methods*, *53*(4), 1563–1580. https://doi.org/10.3758/s13428-020-01500-6

Christensen, A. P., Golino, H., & Silvia, P. J. (2020). A Psychometric Network Perspective on the Validity and Validation of Personality Trait Questionnaires. *European Journal of Personality*, *34*(6), 1095–1108. https://doi.org/10.1002/per.2265

Cohen, S., & Wills, T. A. (1985). Stress, social support, and the buffering hypothesis. *Psychological Bulletin*, *98*(2), 310.

Curtin, N., Malley, J., & Stewart, A. J. (2016). Mentoring the next generation of faculty: Supporting academic career aspirations among doctoral students. *Research in Higher Education*, *57*(6), 714–738.

Demerouti, E., Bakker, A. B., Nachreiner, F., & Schaufeli, W. B. (2001). The job demands-resources model of burnout. *Journal of Applied Psychology*, *86*(3), 499.

Epskamp, S., & Fried, E. I. (2018). A tutorial on regularized partial correlation networks. *Psychological Methods*, *23*(4), 617.

Evans, T. M., Bira, L., Gastelum, J. B., Weiss, L. T., & Vanderford, N. L. (2018). Evidence for a mental health crisis in graduate education. *Nature Biotechnology*, *36*(3), 282–284. https://doi.org/10.1038/nbt.4089

Ferguson, C. L., & McHenry-Sorber, E. (2025). "It's hard not to tie your self-worth to your work:" Faculty Mentorship and Graduate Student Self-Efficacy. *Journal of College Student Retention: Research, Theory & Practice*, 15210251251383061. https://doi.org/10.1177/15210251251383061

Golino, H., & Christensen, A. (2024). EGAnet: Exploratory Graph Analysis–A Framework for Estimating the Number of Dimensions in Multivariate Data Using Network Psychometrics.(2025). doi: 10.32614/CRAN. package. *EGAnet. Accessed*, *20*.

Golino, H. F., & Epskamp, S. (2017). Exploratory graph analysis: A new approach for estimating the number of dimensions in psychological research. *PloS One*, *12*(6), e0174035.

Greenhaus, J. H., & Beutell, N. J. (1985). Sources of Conflict between Work and Family Roles. *The Academy of Management Review*, *10*(1), 76. https://doi.org/10.2307/258214

House, J. S., Landis, K. R., & Umberson, D. (1988). Social Relationships and Health. *Science*, *241*(4865), 540–545. https://doi.org/10.1126/science.3399889

Hyun, S. H., Rogers, J. G., House, S. C., Sorkness, C. A., & Pfund, C. (2022). Revalidation of the Mentoring Competency Assessment to evaluate skills of research mentors: The MCA-21. *Journal of Clinical and Translational Science*, *6*(1), e46.

Jiménez, M., Abad, F. J., Garcia-Garzon, E., Golino, H., Christensen, A. P., & Garrido, L. E. (2025). Dimensionality assessment in bifactor structures with multiple general factors: A network psychometrics approach. *Psychological Methods*, *30*(4), 770.

Kossek, E. E., Pichler, S., Bodner, T., & Hammer, L. B. (2011). Workplace social support and work–family conflict: A meta-analysis clarifying the influence of general and work–family-specific supervisor and organizational support. *Personnel Psychology*, *64*(2), 289–313. https://doi.org/10.1111/j.1744-6570.2011.01211.x

Liu, C., Wang, L., Qi, R., Wang, W., Jia, S., Shang, D., Shao, Y., Yu, M., Zhu, X., Yan, S., Chang, Q., & Zhao, Y. (2019). Prevalence and associated factors of depression and anxiety among doctoral students: The mediating effect of mentoring relationships on the association between research self-efficacy and depression/anxiety. *Psychology Research and Behavior Management*, *Volume 12*, 195–208. https://doi.org/10.2147/PRBM.S195131

Maslach, C. (2011). Burnout and engagement in the workplace: New perspectives. *European Health Psychologist*, *13*(3), 44–47.

Nielson, T. R., Carlson, D. S., & Lankau, M. J. (2001). The supportive mentor as a means of reducing work–family conflict. *Journal of Vocational Behavior*, *59*(3), 364–381.

Pajares, F. (1996). Self-Efficacy Beliefs in Academic Settings. *Review of Educational Research*, *66*(4), 543–578. https://doi.org/10.3102/00346543066004543

Pyhältö, K., Haverinen, K., Vekkaila, J., & Peltonen, J. (2017). Doctoral students' social support profiles and their relationship to burnout, drop-out intentions, and time to candidacy. *International Journal of Doctoral Studies*, *12*, 157–173.

Sitzmann, T., & Yeo, G. (2013). A Meta-Analytic Investigation of the Within-Person Self-Efficacy Domain: Is Self-Efficacy a Product of Past Performance or a Driver of Future Performance? *Personnel Psychology*, *66*(3), 531–568. https://doi.org/10.1111/peps.12035

Tikkanen, L., Ketonen, E., Toom, A., & Pyhältö, K. (2025). PhD candidates' and supervisors' wellbeing and experiences of supervision. *Higher Education*, *90*(5), 1451–1469. https://doi.org/10.1007/s10734-024-01385-w

Tuma, T. T., Fedesco, H. N., Rosenzweig, E. Q., Chen, X.-Y., & Dolan, E. L. (2025). Seeing isn't believing? Mixed effects of a perspective-getting intervention to improve mentoring relationships for science doctoral students. *CBE—Life Sciences Education*, *24*(4), ar44. https://doi.org/10.1187/cbe.25-05-0099
